# High-resolution *in vivo* identification of miRNA targets by Halo-Enhanced Ago2 Pulldown

**DOI:** 10.1101/820548

**Authors:** Xiaoyi Li, Yuri Pritykin, Carla P. Concepcion, Yuheng Lu, Gaspare La Rocca, Minsi Zhang, Peter J. Cook, Yu Wah Au, Olesja Popow, Joao A. Paulo, Hannah G. Otis, Chiara Mastroleo, Paul Ogrodowski, Ryan Schreiner, Kevin M. Haigis, Doron Betel, Christina S. Leslie, Andrea Ventura

**Author notes:** Corresponding authors: Andrea Ventura, Christina S. Leslie. These authors contributed equally to this work (Xiaoyi Li, Yuri Pritykin, Carla P. Concepcion).

## Abstract

The identification of miRNA targets by Ago2 crosslinking-immunoprecipitation (CLIP) methods has provided major insights into the biology of this important class of non-coding RNAs. However, these methods are technically challenging and not easily translated to an *in vivo* setting. To overcome these limitations and to facilitate the investigation of miRNA functions in mice, we have developed a method (HEAP: for Halo-Enhanced Ago2 Pulldown) to map miRNA-mRNA binding sites. This method is based on a novel genetically engineered mouse harboring a conditional, Cre-regulated, Halo-Ago2 allele expressed from the endogenous Ago2 locus. By using a resin conjugated to the HaloTag ligand, Ago2-miRNA-mRNA complexes can be efficiently purified from cells and tissues expressing the endogenous Halo-Ago2 allele. We demonstrate the reproducibility and sensitivity of this method in mouse embryonic stem cells, in developing embryos, in adult tissues and in autochthonous mouse models of human brain and lung cancers.

The tools and the datasets we have generated will serve as a valuable resource to the scientific community and will facilitate the characterization of miRNA functions under physiological and pathological conditions.

## INTRODUCTION

A key challenge in deciphering the biological functions of miRNAs remains the identification of their targets *in vivo* under physiologic and pathologic conditions. Although significant progress has been made in computational methods to predict miRNA binding sites (Agarwal et al., 2015; Bartel, 2009; Grimson et al., 2007), these methods do not take into account the many known and unknown variables that determine whether a ‘potential’ target site is in fact available and bound by a miRNA in a given cellular context. To complement computational approaches, biochemical methods to purify Ago2-miRNA-mRNA complexes have been developed (Chi et al., 2009; Hafner et al., 2010; Helwak et al., 2013; Konig et al., 2010; Moore et al., 2015; Van Nostrand et al., 2016). Although the details vary, these methods all rely on the use of antibodies to precipitate Argonaute-containing complexes, usually after UV crosslinking, followed by high-throughput sequencing of the associated mRNAs.

While these methods have been applied with substantial success to map miRNA-mRNA interactions in cell lines, they are used much less extensively *in vivo* due to their technical complexity and the lack of efficient ways to restrict the analysis to specific cell types within a tissue. To overcome these limitations, we have developed a novel method Halo-enhanced Ago2 pulldown (HEAP), which utilized a tagged version of the Ago2 protein and allows the direct purification of Ago2-containing complexes bypassing the need for radiolabeling, immunoprecipitation, and gel purification. To facilitate the application of this method *in vivo*, we have generated a novel mouse strain in which a conditional allele of Halo-tagged Ago2 is knocked into the endogenous Ago2 locus and is activated upon exposure to Cre recombinase.

To benchmark the HEAP method, we applied it to identify miRNA targets in diverse cellular contexts, including murine embryonic stem cells (mESCs), wild type and miR-17∼92-null mid-gestation mouse embryos, adult mouse lungs, adult mouse brains, and three distinct autochthonous mouse models of human lung and brain cancers. As a result, we have identified a large number of miRNA targets at high resolution and demonstrated the reproducibility and sensitivity of the HEAP method.

The datasets and the tools generated in this study reveal the complex landscape of miRNA targeting *in vivo* and will facilitate future studies aimed at characterizing the biological functions of this important class of small non-coding RNAs under physiological and pathological conditions.

## RESULTS

### A Halo-Ago2 fusion protein enables antibody-free purification of miRNA targets

The HaloTag is a 33 kDa haloalkane dehalogenase encoded by the DhaA gene from *Rhodococcus rhodochrous* that has been mutagenized to form an irreversible covalent bond to synthetic chloroalkane ligands (collectively known as HaloTag ligands) (Encell et al., 2012; Los et al., 2008). Linking the chloroalkane ligand to a solid substrate enables the efficient purification of fusion proteins containing the HaloTag (**Figure 1a**).

**Figure 1.**
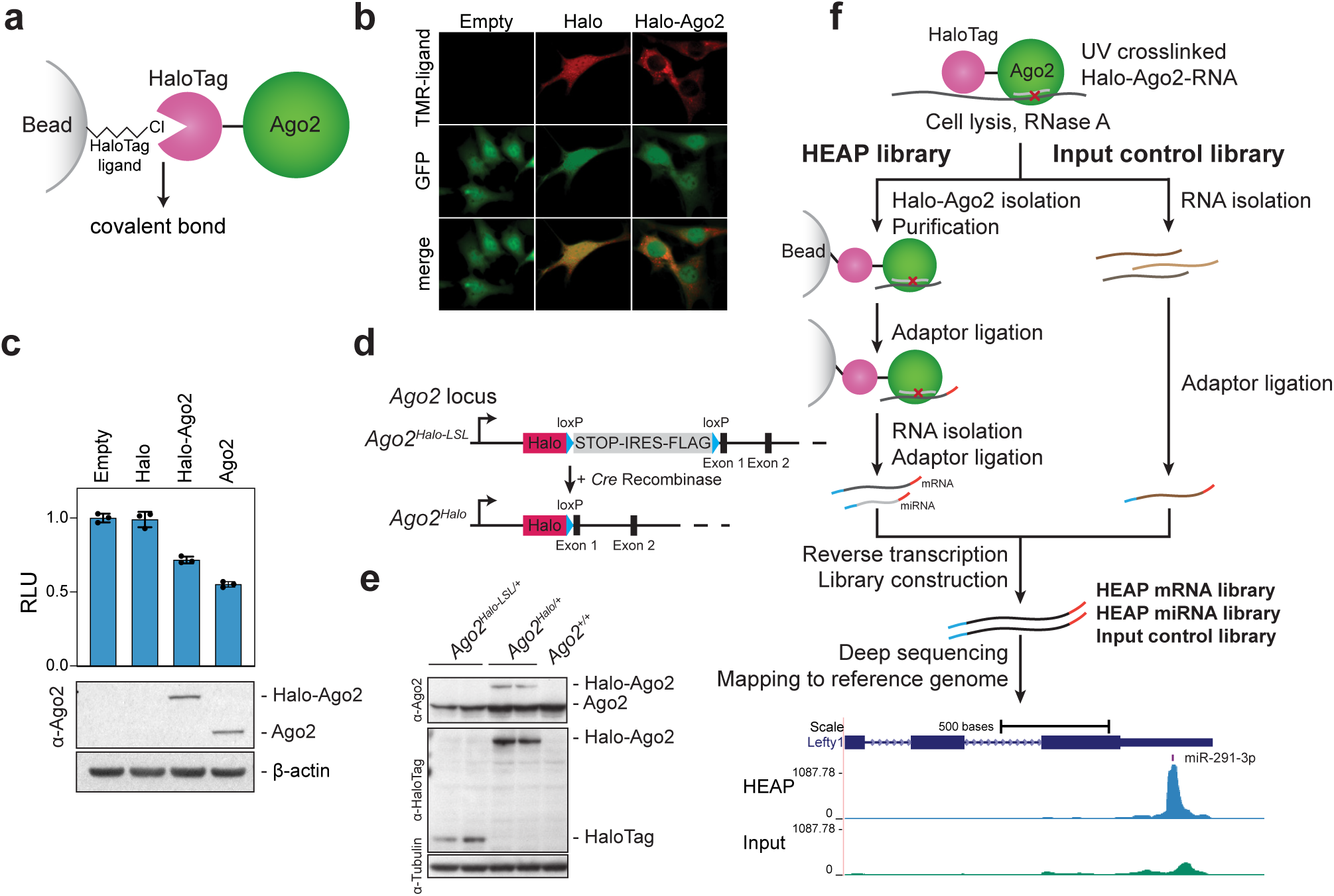
Halo-enhanced Ago2 pulldown (HEAP). **a)** Schematic of the Halo-Ago2 fusion protein covalently bound to a bead-conjugated Halo-ligand. **b)** Ago2-null immortalized MEFs transduced with retroviruses MSCV-PIG, MSCV-PIG-Halo or MSCV-PIG-Halo-Ago2 were incubated with the HaloTag TMR ligand and imaged. Notice the prevalently cytoplasmic localization of the Halo-Ago2 fusion protein. TMR: Tetramethylrhodamine. **c)** Ago2-null MEFs transduced with retroviral vectors encoding HaloTag alone, full length Ago2 or the Halo-Ago2 fusion protein were transiently transfected with reporter plasmids expressing firefly and renilla luciferase and a plasmid expressing an shRNA against the firefly luciferase. The ratio between firefly and renilla luciferase activity was measured 48 hours after transfection (upper panel). Whole-cel lysates of the MEFs were probed with antibodies against Ago2 and β-actin (lower panel). **d)** Schematic of the targeting strategy used to generate the Halo-Ago2 conditional knock-in allele. Halo: HaloTag; STOP: stop codon; IRES: internal ribosome entry site. **e)** Whole-cell lysates from mESCs with the indicated genotypes were probed with antibodies against Ago2, HaloTag and Tubulin. **f)** Outline of the strategy used to generate HEAP and input control libraries (upper panel) and a representative Halo-Ago2 binding site identified in mESCs (lower panel).

To determine whether a HaloTag-based strategy can be employed to purify complexes containing Ago2 proteins bound to miRNA and target mRNAs, we fused the HaloTag to the N-terminus of Ago2 (Halo-Ago2; **Figure 1a**). When expressed in Ago2-null mouse embryonic fibroblasts (MEFs, O’Carroll et al., 2007), the Halo-Ago2 fusion protein localized largely to the cytoplasm, while the HaloTag alone displayed uniform localization to both the cytoplasm and the nucleus (**Figure 1b**). Importantly, the Halo-Ago2 construct was nearly as effective as wild type Ago2 at rescuing RNAi in Ago2-null MEFs, indicating that the Halo-Ago2 fusion protein retains slicing activity (**Figure 1c**).

To avoid artifacts due to ectopic expression of Halo-Ago2 and to enable the isolation of Ago2 complexes directly from murine tissues, we inserted the HaloTag cassette in the endogenous Ago2 locus in mESCs (**Figure 1d and Supplementary Figure 1**). In this knock-in allele, the HaloTag is separated from the first coding exon of Ago2 by an in frame loxP-STOP-IRES-FLAG-loxP (LSL) cassette (Ago2^Halo-LSL^). Cells harboring this allele express a bicistronic mRNA encoding for two proteins: the HaloTag and a Flag-Ago2 fusion protein whose translation is initiated by an internal ribosomal entry site (IRES). Upon expression of the Cre recombinase, the LSL cassette is excised and the HaloTag is now brought in frame with the first coding exon of Ago2, thus resulting in the expression of the Halo-Ago2 fusion protein (**Figure 1d-e and Supplementary Figure 1**). The recombined allele expressing the Halo-Ago2 fusion will be hereafter referred to as Ago2^Halo^.

We first tested whether the Ago2^Halo^ allele could be used to map miRNA-mRNA interactions in mESCs. For these experiments, we adapted the HITS-CLIP method originally developed by the Darnell group (Chi et al., 2009) with two significant streamlining modifications enabled by the covalent bond between Halo-Ago2 and the HaloTag ligand. First, instead of using anti-Ago2 antibodies to isolate Ago2-containing complexes, we used sepharose beads covalently linked to the HaloTag ligand. Second, the radiolabeling and SDS-PAGE purification step necessary in CLIP protocols to purify RNAs bound to Ago2 were omitted and replaced by extensive washes followed by direct RNA extraction from beads, library construction, and high-throughput sequencing of Halo-Ago2-bound miRNA and mRNAs (**Figure 1f**). We refer to this method as Halo-enhanced Ago2 pulldown (HEAP). By performing HEAP, two types of libraries are generated: a target library (mRNAs) and a miRNA library (**Figure 1f, Supplementary Figure 2a**). The former allows the identification of miRNA binding sites on their targets, while the latter provides an estimate of miRNA abundance.

When mapped to the mouse genome, HEAP mRNA libraries generated from Ago2^Halo/+^ mESCs—but not those generated from control Ago2^Halo-LSL/+^ cells—produced well-defined “clusters” of reads, from here on referred to as peaks (**Supplementary Figure 2a-b**). To facilitate the identification of these peaks, we adapted the “SMinput” protocol used in eCLIP (Van Nostrand et al., 2016) and generated input control libraries from size-matched RNA fragments after the limited RNase protection step (**Figure 1f**). We first identified putative peaks using the CLIPanalyze package (https://bitbucket.org/leslielab/clipanalyze), an improved peak-calling algorithm based on edge detection technique similar to methods from image processing (Hsin et al., 2018; Lianoglou et al., 2013; Loeb et al., 2012). Then, CLIPanalyze utilized the input control libraries as background to assign *p*-value to each peak, using library size normalization based on reads aligned across the genome outside of putative peaks (see Methods for additional details).

Previous studies have demonstrated that 3’-untranslated regions (3’UTRs) of mRNAs are the preferred, although not exclusive, sites of interaction between miRNAs and mRNAs (Bartel, 2018; Chi et al., 2009; Sarshad et al., 2018). Consistent with these findings, the majority of HEAP peaks we identified in mESCs mapped to 3’UTRs, followed by sites mapping to coding exons (**Figure 2a, Supplementary Figure 2c**). The fraction of 3’UTR and exonic peaks increased monotonically with their statistical significance, while intergenic and intronic peaks had the opposite behavior. For example, when examining the 1,000 most statistically significant peaks, greater than 50% of them mapped to 3’UTRs and less than 3% mapped to introns (**Figure 2a**). To measure reproducibility, we performed pairwise Irreproducible Discovery Rate (IDR) (Li et al., 2011) analysis on three independent HEAP libraries generated from mESCs. On average, this analysis identified 80% of peaks as reproducible at IDR < 0.05, demonstrating the robustness of the HEAP method (**Supplementary Figure 2b, d**).

**Figure 2.**
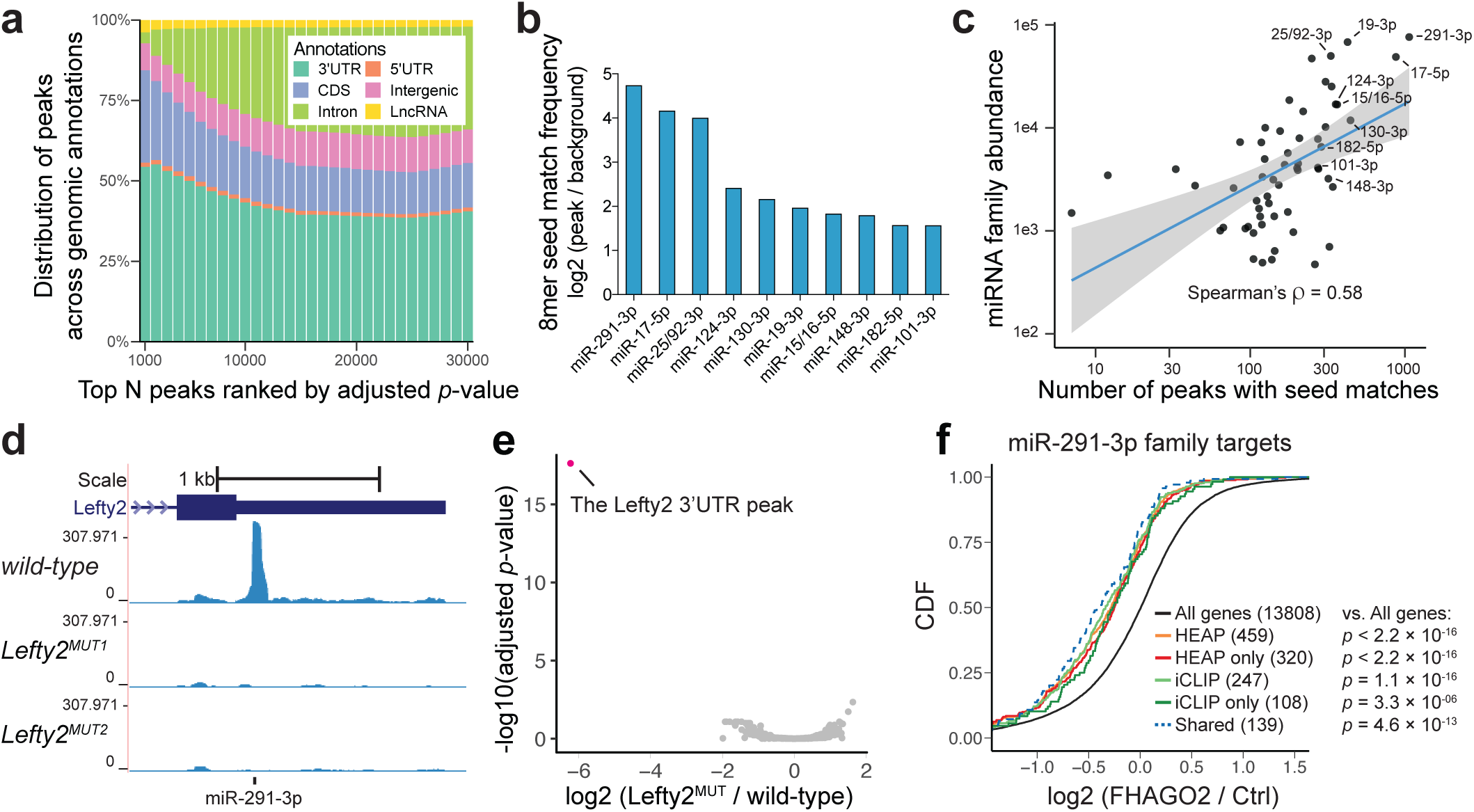
Mapping Halo-Ago2 binding sites in mESCs. **a)** Peaks identified in the HEAP libraries from mESCs were ranked by increasing adjusted *p*-value before calculating their distribution across genomic annotations. CDS: coding sequence; 5’UTR: 5’-untranslated region; 3’UTR: 3’-untranslated region. LncRNA: long non-coding RNA. **b)** Enrichment for sequences complementary to murine miRNA seeds (8mers) was calculated comparing 3’UTR sequences within and outside HEAP peaks. The bar plot shows enrichment for the top ten seed matches. **c)** Scatter plot showing the correlation between number of 3’UTR peaks with 7mer or 8mer seed matches to individual miRNA families and abundance of their corresponding miRNAs as measured in HEAP miRNA libraries. Blue line: best fit linear regression, with 95% confidence interval in grey. **d)** Genome browser view of the Lefty2 3’UTR with tracks corresponding to libraries generated from wild-type mESC (wild-type) or cells harboring targeted mutations disrupting the predicted miR-291-3p binding site (Lefty2^MUT1^, Lefty2^MUT2^). **e)** Volcano plot of global changes in HEAP peak intensity between the Lefty2^MUT^ and wild-type mESC libraries. Notice the selective loss of the Lefty2 3’UTR binding site (highlighted). **f)** Cumulative distribution function (CDF) plot for targets of miR-291-3p identified by iCLIP or HEAP. The log2 fold change was calculated in Ago1-4^-/-^ mESCs upon ectopic FHAGO2 expression (dataset from Bosson et al., 2014; GSE61348). HEAP only: targets identified uniquely by HEAP; iCLIP only: targets identified uniquely by iCLIP; shared: targets identified by both methods (*p*-value: one-sided Kolmogorov–Smirnov test).

To gain additional insights into the nature of peaks identified by HEAP, we ran an unbiased 7-nt sequence (7-mer) enrichment analysis on the sequences underlying peaks mapped to 3’UTRs (**Supplementary Figure 3a**). Inspection of the resulting motifs revealed a marked enrichment for seed matches corresponding to miRNA families whose members are collectively highly expressed in mESCs (**Supplementary Figure 3b, Figure 2b**). We observed a positive correlation between the relative abundance of individual miRNA families—estimated from the miRNA libraries—and the number of corresponding peaks identified by HEAP (Spearman’s ρ = 0.58) (**Figure 2c**).

To directly test whether the peaks identified by HEAP reflect true miRNA-mRNA interactions, we selected a robust peak identified in the 3’UTR of the Lefty2 mRNA (**Figure 2d**). The sequence underlying this peak includes a highly conserved 8-mer that is complementary to the miR-291-3p seed (**Supplementary Figure 4a**). We used CRISPR-Cas9 and homologous recombination in mESCs to introduce point mutations designed to disrupt this seed match (**Supplementary Figure 4b**). HEAP libraries generated from two independent clones thus generated showed complete and selective loss of the Lefty2 peak, further demonstrating the ability of the HEAP method to map *bona fide* miRNA-mRNA interactions in cells (**Figure 2d-e**).

To assess the ability of HEAP to identify functional miRNA binding sites, we analyzed an RNA-seq dataset generated by Bosson and colleagues from mESCs null for all four Argonaute proteins (Ago1-4^-/-^) in the presence or absence of exogenously expressed Flag- and HA-tagged AGO2 (FHAGO2) [(Bosson et al., 2014), GSE61348]. Introduction of FHAGO2 in Ago1-4^-/-^ cells should restore miRNA function, inducing repression of miRNA targets. In agreement with this prediction, miRNA targets identified by HEAP were preferentially repressed upon FHAGO2 reintroduction, as illustrated by cumulative distribution function (CDF) plots (**Supplementary Figure 5a**). Furthermore, the effect was stronger for targets assigned by HEAP to the most abundantly expressed miRNA families in mESCs, in line with the hypothesis that miRNA-target stoichiometry is a major determinant of gene repression. For example, we observed the strongest repression for targets of the miR-291-3p, miR-17-5p and miR-148-3p families, three miRNA families that alone account for greater than 12% of all miRNAs in mESCs (**Supplementary Figure 5a and data not shown**).

This analysis also allowed us to compare miRNA targets identified by HEAP to those previously identified by Bosson et al. in Ago1-4^-/-^-FHAGO2 mESCs using iCLIP, a well-established variant of HITS-CLIP (Konig et al., 2010). By applying the CLIPanalyze algorithm, we identified 6,813 FHAGO2 binding sites in the iCLIP library generated from Ago1-4^-/-^-FHAGO2 mESCs, and nearly twice as many (on average 13,532) in each of the three HEAP mESC libraries (*p*-value < 0.1). Additionally, the iCLIP library displayed significantly fewer peaks mapping to 3’UTR and more peaks mapping to intergenic regions compared to the HEAP libraries (**Supplementary Figure 5b**).

3’UTR targets for miR-291-3p seed family identified by both methods were associated with strong repression of the corresponding genes upon FHAGO2 reintroduction (**Figure 2f**). The overlap between miR-291-3p binding sites identified by iCLIP and HEAP in 3’UTRs was partial, with the HEAP target pool being nearly twice as large (**Supplementary Figure 5c**). Importantly, the targets identified only by HEAP also displayed strong repression upon FHAGO2 reintroduction, indicating that they are functional miRNA binding sites (**Figure 2f**).

We further confirmed the ability of HEAP to identify functional miRNA targets by measuring mRNA and protein expression changes of HEAP targets upon inactivation of Dicer1, the key enzyme responsible for miRNA maturation, in mESCs (**Supplementary Figure 6**).

Collectively, these results demonstrated that the HEAP method provides a simple, effective and sensitive method to identify miRNA-mRNA interactions in cells.

### A conditional Halo-Ago2 mouse enables identification of miRNA-mRNA interactions *in vivo*

The accurate identification of miRNA targets in vivo and in a cell-type specific context is essential to dissect the functions of miRNAs in development, homeostasis, and disease. To translate the HEAP method to an in vivo setting, we used mESCs harboring the Cre-inducible Halo-Ago2 allele to generate Ago2Halo-LSL/+ mice. We then crossed these animals to CAGGS-Cre mice (Araki et al., 1997) to delete the LSL cassette and induce ubiquitous expression of the endogenous Halo-Ago2 allele. PCR and immunoblot analysis in mouse embryonic fibroblasts (MEFs) and tissues derived from these mice confirmed efficient deletion of the LSL cassette and expression of the Halo-Ago2 protein (Supplementary Figure 7a-b and Figure 3a). Although Ago2Halo/+ and Ago2Halo-LSL/+ mice were obtained at the expected Mendelian frequency and were phenotypically indistinguishable from wild-type mice, homozygous mice for the Ago2Halo or the Ago2Halo-LSL alleles were recovered at sub-Mendelian frequencies (9.9% and 11.9%, respectively, compared to the expected 25%, Figure 3b). The reduced viability of homozygous mutant mice might reflect the reduced expression of the Halo-Ago2 allele compared to wild-type Ago2 (Figure 3a and Supplementary Figure 7b). Although we cannot exclude the possibility that the presence of the HaloTag subtly interferes with assembly or function of the miRNA-induced silencing complex (miRISC), size-exclusion chromatography experiments indicated that the Halo-Ago2 fusion protein co-eluted with endogenous Ago2 in high molecular weight complexes (Supplementary Figure 7c) and physically interacts with Tnrc6a, one member of the Tnrc6 family of scaffold proteins responsible for the assembly of the miRISC (**Supplementary Figure 7d**).

**Figure 3.**
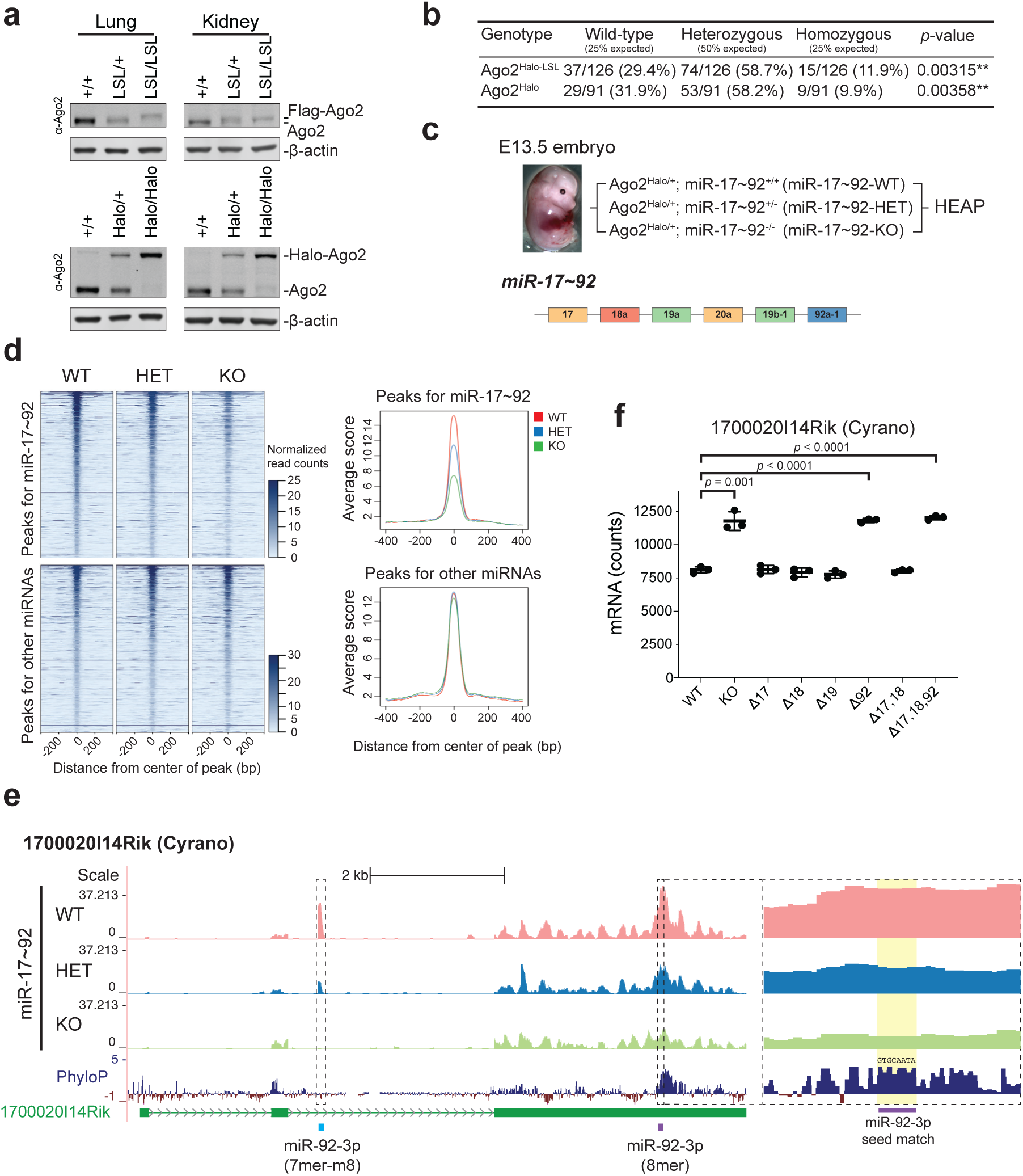
Identification of miR-17∼92 targets in E13.5 embryos. **a)** Expression of Ago2 fusion proteins in the lungs and kidneys of Ago2Halo-LSL and Ago2Halo mice. +/+: wild-type; LSL: Ago2Halo-LSL; Halo: Ago2Halo. **b)** Absolute numbers and frequencies of genotypes obtained from heterozygous intercrosses of Ago2Halo-LSL/+ or Ago2Halo/+ mice (*p*-value: Chi-Squared test). **c)** Outline of the HEAP experiments in E13.5 Ago2Halo/+ embryos wild-type, heterozygous or homozygous knockout for the miR-17∼92 cluster. Short names for the genotypes were created. A schematic of the miR-17∼92 cluster is shown at the bottom. miRNA members are colored based on their seed families. **d)** Heatmap and histogram of read counts in an 800-bp region surrouding HEAP peaks obtained from miR-17∼92-WT, miR-17∼92-HET and miR-17∼92-KO E13.5 embryos. Peaks containing seed matches for the top 31 miRNA seed families in the miR-17∼92-WT embryo ranked by abundance were chosen. Peaks with seed matches for miRNAs belonging to the miR-17∼92 cluster are plotted in the upper panels, while the remaining peaks are plotted in the lower panels. **e)** Genome browser view of the miR-92a-1-dependent miRNA binding sites detected in the long non-coding RNA 1700020I14Rik (Cyrano). PhyloP: placental mammal basewise conservation by PhyloP. Notice the highly conserved 8mer seed match for miR-92-3p under the second peak. **f)** mRNA expression of Cyrano in the heart of E9.5 embryos harboring an allelic series of miR-17∼92 mutant alleles (RNA-seq data obtained from Han et al., 2015; GSE63813). KO: embryos null for the entire miR-17∼92 cluster; Δ17: embryos null for miR-17 and miR-20a; Δ18: embryos null for miR-18a; Δ19: embryos null for miR-19a and miR-19b-1, Δ 92: embryos null for miR-92a-1; Δ17,18: embryos null for miR-17, miR-18a and miR-20a; Δ17,18,92: embryos null for miR-17, miR-18a, miR-20a and miR-92a-1. Notice that Cyrano is only up-regulated in mutants in which miR-92a-1 is deleted (*p*-value: unpaired t test).

To test whether endogenously expressed Halo-Ago2 can be used to identify miRNA targets *in vivo*, we crossed Ago2^Halo/+^ mice to mice harboring a targeted deletion of the miR-17∼92 locus, a polycistronic miRNA cluster encoding six distinct miRNAs, which has been shown to be essential for mammalian development (Han et al., 2015; Ventura et al., 2008). We then generated HEAP libraries from Ago2^Halo/+^; miR-17∼92^+/+^ (miR-17∼92-WT), Ago2^Halo/+^; miR-17∼92^+/-^ (miR-17∼92-HET) and Ago2^Halo/+^; miR-17∼92^-/-^ (miR-17∼92-KO) E13.5 embryos (**Figure 3c)**. At an adjusted *p*-value cutoff of 0.01, HEAP identified a total of 8,661 peaks in these libraries, with a distribution across genomic annotations similar to that observed in mESCs (**Supplementary Figure 8a**). Importantly, the intensity of peaks overlapping with potential seed matches to members of the miR-17∼92 cluster was greatly reduced—in a dose-dependent fashion—in the libraries generated from miR-17∼92-HET and miR-17∼92-KO embryos (**Fig. 3d and Supplementary Figure 8b-c**). The murine genome contains two additional miRNA clusters that are paralogs to miR-17∼92 and encode similar miRNAs (Ventura et al., 2008), which may explain some residual Halo-Ago2 binding to these sites even in the homozygous mutants. Using a published RNA-seq dataset previously generated in the lab from E9.5 embryos harboring an allelic series of miR-17∼92 mutant alleles [(Han et al., 2015), GSE63813], we demonstrated that HEAP targets containing seed matches for miR-17/20-5p, miR-19-3p and miR-92-3p mediated strong target repression (**Supplementary Figure 9**). Interestingly, we also identified a sizeable fraction of reproducible peaks (4%) mapping to non-coding RNAs. These included two previously uncharacterized miR-17∼92-dependent sites matching the miR-92-3p seed in the long non-coding RNA Cyrano (Kleaveland et al., 2018; Ulitsky et al., 2011) **(Figure 3e)**. Supporting the hypothesis that these peaks are functionally relevant, we observed upregulation of Cyrano in mouse E9.5 embryos lacking miR-92a-1, but not in mice harboring selective deletion of the other members of the cluster (**Figure 3f)** (Han et al., 2015). These results demonstrate the usefulness of the Halo-Ago2 mouse strain in facilitating the identification of miRNA targets *in vivo* and suggest that additional studies aimed at determining the functional consequences of loss of miR-92a-1 on *Cyrano* function may be warranted.

To directly compare the performance of HEAP to immunoprecipitation-based approaches *in vivo*, we next generated libraries from the cortex of P13 Ago2^Halo/+^ mice, a tissue from which high-quality miRNA-target libraries have been previously generated by HITS-CLIP and CLEAR-CLIP (Chi et al., 2009; Moore et al., 2015). Two HEAP libraries generated from the cortices of Ago^Halo/+^ mice produced 7,069 peaks at an adjusted *p*-value cutoff of 0.05. This number of miRNA-mRNA interaction sites is comparable to that identified by Moore and colleagues (CLEAR-CLIP, GSE73059, n = 7,927) using 12 biological replicates (**Supplementary Figure 10a-b**). HEAP and CLEAR-CLIP identified similar numbers of targets for miR-124-3p, one of the most abundance miRNAs in the mouse cortex (**Supplementary Figure 10c**). When benchmarked against a microarray gene expression dataset generated from neuroblastoma cells (CAD) ectopically expressing miR-124 [(Makeyev et al., 2007), GSE8498], HEAP and CLEAR-CLIP were equally effective at identifying miR-124 target sites that mediated target repression (**Supplementary Figure 10d**). Collectively, these results demonstrate that the HEAP method provides a simple and cost-effective approach to identify miRNA-mRNA interactions during murine development and in adult tissues.

### Identification of miRNA targets in normal adult tissues and in autochthonous tumors

We next tested whether the conditional Halo-Ago2 mouse could be used to identify miRNA-mRNA interactions in primary autochthonous tumors and in their tissues of origin. We first chose a mouse model of glioma driven by the *Bcan-Ntrk1* gene fusion that we was recently developed in our laboratory (Cook et al., 2017). In this model, p53^fl/fl^ mice are injected intracranially with a mixture of two recombinant adenoviruses. The first expresses Cas9 and two sgRNAs (Ad-BN) designed to induce the *Bcan-Ntrk1* rearrangement, an intra-chromosomal deletion resulting in the fusion between the N-terminal portion of Bcan and the kinase domain of Ntrk1. The second adenovirus expresses the Cre recombinase (Ad-Cre) to achieve concomitant deletion of p53 and allow glioma formation. By performing this procedure in 4∼6-week-old p53^fl/fl^; Ago2^Halo-LSL/+^ mice, we produced Bcan-Ntrk1 driven gliomas expressing the endogenous Halo-Ago2 allele.

We generated HEAP libraries from three independent Bcan-Ntrk1 gliomas and from the normal cortices of three age-matched Ago2^Halo/+^ mice. Quantification of miRNA abundance in HEAP miRNA libraries revealed drastic differences between the two tissues, with 77 miRNA seed families (26 broadly conserved) being significantly upregulated in gliomas, and 77 families (18 broadly conserved) downregulated (adjusted *p*-value < 0.05, absolute log2FC > 0.5, **Figure 4a**). Of note, the significantly downregulated families include miR-124-3p and miR-128-3p, two miRNA families that are highly expressed in the cortex of mice (Bak et al., 2008; Landgraf et al., 2007) and functionally important in the mouse central nervous system as suggested by genetic loss-of-function studies (Sanuki et al., 2011; Tan et al., 2013). Additionally, members of the oncogenic miRNA cluster miR-17∼92 (He et al., 2005; Ota et al., 2004) were among the most strongly upregulated miRNAs in gliomas, suggesting the possibility that these miRNAs are functionally relevant in gliomagenesis.

**Figure 4.**
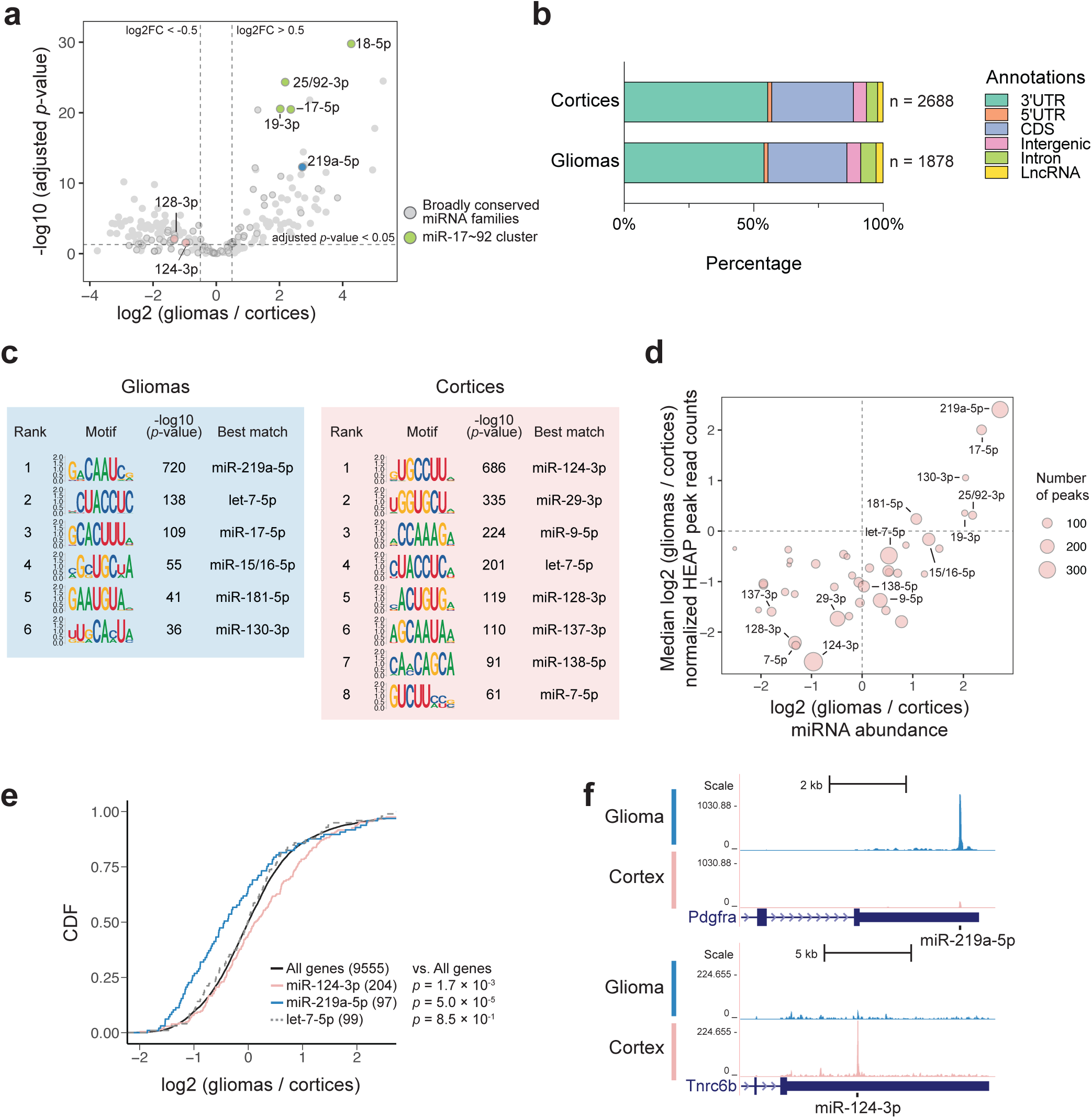
Mapping miRNA-target interactions in normal brain and brain tumors. **a)** Volcano plot of global changes in miRNA family expression between normal cortices and Bcan-Ntrk1-driven gliomas, as determined by HEAP. Broadly conserved families are highlighted in circles and a few miRNA families of interest are colored and annotated. **b)** Total number and distribution across genomic annotations of peaks identified in the cortex and glioma HEAP libraries at adjusted *p*-value < 0.05. **c)** Top differentially enriched 8-mers identified in glioma and cortex HEAP peaks (cutoff: adjusted *p*-value < 0.05; absolute log2 (gliomas / cortices) > 0.5) by the HOMER *de novo* motif discovery algorithm. miRNA families whose seed sequences are complementary to these motifs are annotated. **d)** Changes in peak intensity correlate with changes in miRNA abundance. The area of each circle is proportional to the number of targets of each miRNA seed family as identified by HEAP. Only broadly conserved miRNA families with more than 40 HEAP targets are shown. **e)** Cumulative distribution function (CDF) plot for targets of miR-124-3p,miR-219a-5p and let-7-5p identified by HEAP. mRNA expressions were estimated using read counts in input control libraries. The mRNA log2 fold change was calculated as gliomas vs. cortices (*p*-value: one-sided Kolmogorov–Smirnov test). **f)** Genome browser view of representative Halo-Ago2 binding sites detected exclusively in gliomas (top) or cortices (bottom).

Using an adjusted *p*-value cutoff 0.05, we identified 1,878 Halo-Ago2 binding sites in tumors and 2,688 sites in normal cortices, with an overlap of 1,335 sites. Peaks distribution across genomic annotations was similar between the two tissues, with the majority of peaks mapping to 3’UTRs (**Figure 4b)**. On average, 60% of peaks from each biological replicate of glioma libraries passed an IDR cutoff of 0.05 (**Supplementary Figure 11**).

Analysis of seed matches under the peaks revealed marked differences between normal and neoplastic brains. Motifs complementary to the seeds of miR-219a-5p, miR-17-5p, miR-15/16-5p, miR-181-5p and miR-130-3p were preferentially enriched in peaks identified in gliomas, while motifs complementary to the seeds of miR-124-3p, miR-29-3p, miR-9-5p, miR-128-3p, miR-137-3p, miR-138-5p and miR-7-5p were preferentially enriched in peaks from normal cortices (adjusted *p*-value < 0.1, absolute log2FC (gliomas vs. cortices) > 0.5). Targets for the let-7-5p family of miRNAs were also abundant, but not differentially represented between the normal brain and tumors (**Figure 4c, f and Supplementary Figure 12**). The enrichment for specific seed matches observed in the two conditions reflected in large part the differential expression of the corresponding miRNAs (**Figure 4d**) and resulted in differential gene regulation, as demonstrated by a statistically significant repression of miR-219a-5p targets in gliomas and of miR-124-3p targets in the normal cortices (**Figure 4e**).

Among all miRNA families, the miR-219a-5p family had the highest number of targets in gliomas (300 out of 1,878 peaks contained 6mer, 7mer or 8mer seed matches to miR-219a-5p). miR-219-5p has been reported to regulate oligodendrocyte (OL) differentiation and myelination in mice via targeting important regulators of oligodendrocyte progenitor cell (OPC) maintenance (Dugas et al., 2010; Emery, 2010; Fan et al., 2017; Wang et al., 2017; Zhao et al., 2010). Interestingly, we observed a strong interaction between miR-219a-5p and Pdgfra (**Figure 4f**), a characteristic marker of OPCs and a key player in gliomagenesis.

Finally, to extend the application of the HEAP method to other tumor types, we mapped miRNA-mRNA interactions in two murine models of non-small cell lung cancer (NSCLC): the Cre recombinase-mediated KRas^LSL-G12D/+^; p53^fl/fl^ (KP) model (Jackson et al., 2001) and a CRISPR-Cas9 induced model driven by a chromosomal inversion resulting in the formation of the Eml4-Alk (EA) gene fusion (Maddalo et al., 2014). These two mouse models recapitulate two types of NSCLC observed in humans and differ not only for the initiating genetic lesions but also for the modality with which tumor formation is induced. We generated HEAP libraries from Ago2^Halo-LSL/+^ mice bearing primary KP (N = 2) and EA (N = 3) tumors. Tumor-specific expression of the Halo-Ago2 allele was induced at the time of tumor initiation by intratracheal delivery of Ad-Cre, alone for the KP model or in combination with recombinant adenoviruses expressing Cas9 and the two gRNAs necessary to induce the Eml4-Alk rearrangement in the EA model (Ad-EA). In parallel, we also generated HEAP libraries from the lungs of two Ago2^Halo/+^ mice (**Figure 5a**).

**Figure 5.**
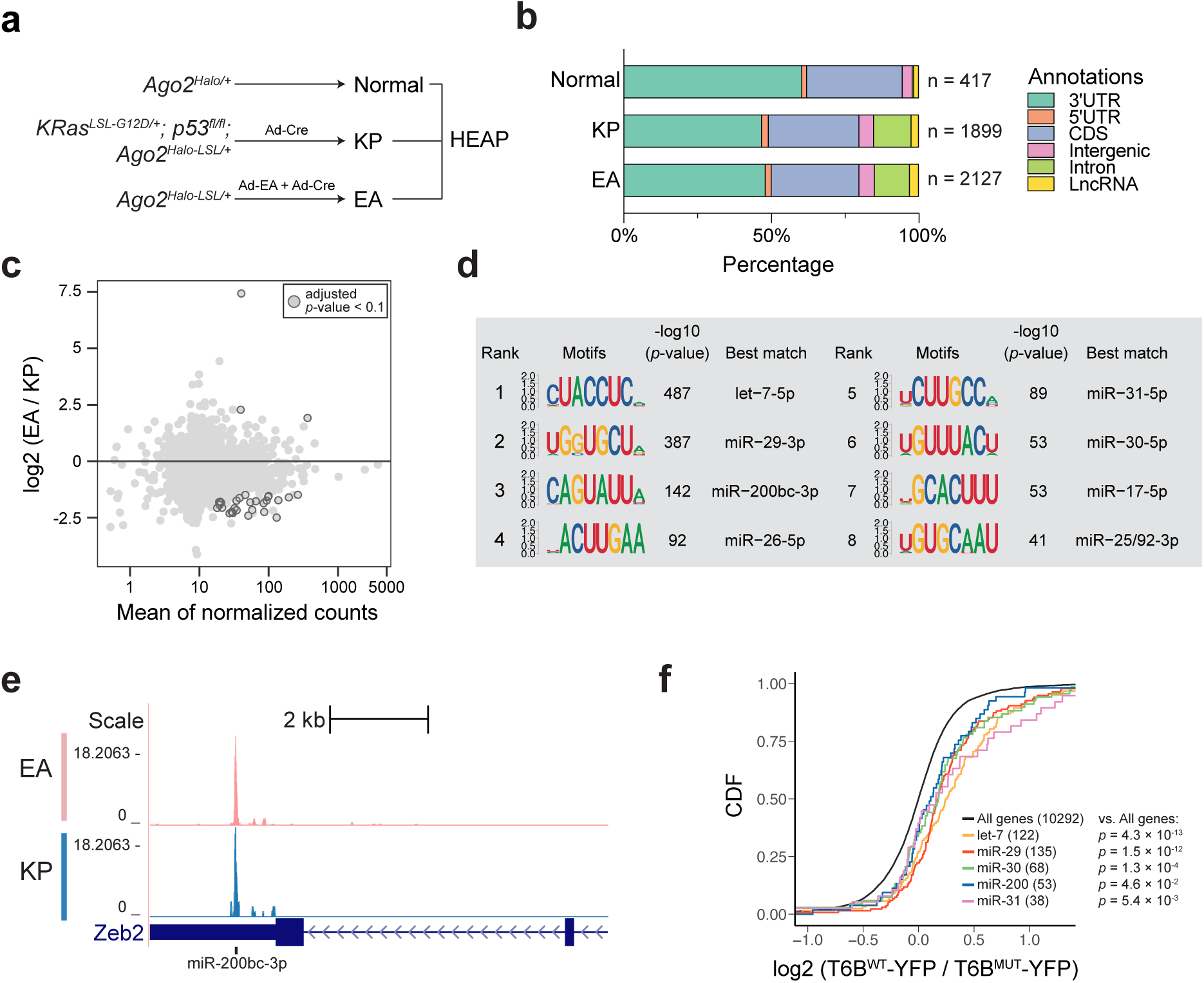
Mapping miRNA-target interactions in lung adenocarcinomas. **a)** Schematic of the experimental design. **b)**Total number and distribution across genomic annotations of peaks identified in normal lungs (two replicates) and in the KP and EA lung adenocarcinomas (two and three replicates, respectively) at adjusted *p*-value < 0.05. **c)** HEAP peaks are highly similar between KP and EA lung adenocarcinomas. MA plot for peak intensity changes between the KP and EA lung adenocarcinomas is shown. Peaks with adjusted *p*-value lower than 0.1 are highlighted. **d)** Sequence logos of the most enriched 8-mers as determined by the HOMER *de novo* motif discovery algorithm under the HEAP peaks identified specifically in lung adenocarcinomas (cutoff: adjusted *p*-value < 0.05; log2 (tumors / lungs) > 0.5). miRNA families whose seeds are complementary to the motifs are annotated. **e)** Genome browser view of the miR-200bc-3p binding site in the 3’UTR of Zeb2 in both KP and EA HEAP libraries. **f)** Targets identified by HEAP show preferential de-repression upon inactivation of the miRISC. Cumulative distribution function (CDF) plot of mRNA expression changes induced by T6B-YFP expression in a mouse KP lung cancer cell line (T6B^WT^-YFP / T6B^MUT^-YFP). Targets identified by HEAP for the indicated miRNA families were compared to background (“all genes”). *P*-values were determined by one-sided Kolmogorov–Smirnov tests.

The tumor libraries produced 1,899 peaks for the KP tumors and 2,127 peaks for the EA tumors. In contrast, only 417 peaks were identified in normal lungs (**Figure 5b**). This difference could not be attributed to differences in sequencing depth or Halo-Ago2 expression levels in normal lungs vs. tumors (**Supplementary Figure 13a**). Rather, it may reflect reduced levels of fully assembled miRISC in the normal lung compared to lung tumors [(La Rocca et al., 2015) and La Rocca et al., manuscript in preparation].

Surprisingly, a direct comparison of the peaks identified in KP and EA tumors revealed strong correlation between the two tumor types (**Supplementary Figure 13b and Figure 5c**), suggesting that the miRNA targeting landscape is largely independent from the cancer initiation events in these two NSCLC models. Unbiased k-mer frequency analysis visualized as motif enrichment identified distinct miRNA seed-matches enriched in peaks in normal lung and tumors. Binding sites for let-7-5p, miR-29-3p and miR-30-5p were strongly enriched in both tissues, while seed matches for several miRNAs implicated in tumorigenesis and metastasis, such as miR-200bc-3p (Davalos et al., 2012; Gibbons et al., 2009; Gregory et al., 2008; Sato et al., 2017; Si et al., 2017), miR-31-5p (Edmonds et al., 2016), miR-17-5p and miR-25/92-3p (He et al., 2005; Ota et al., 2004) were dominant in the tumor libraries (**Figure 5d and Supplementary Figure 13c**). In human lung adenocarcinomas, miR-200 levels negatively correlate with tumor metastatic potential, at least in part because this miRNA can potently suppress epithelial-to-mesenchymal transition (EMT) (Davalos et al., 2012; Gibbons et al., 2009; Si et al., 2017). In agreement with this model, we observed a strong miR-200bc-3p binding site in the 3’UTR of Zeb2, a master regulator of EMT (**Figure 5e**).

To further validate the functional significance of these miRNA-mRNA interactions in lung cancer, we took advantage of a fusion protein (T6B-YFP) previously shown to bind Argonaute proteins and disrupt assembly of the miRISC complex, leading to a global de-repression of miRNA targets [(Hauptmann et al., 2015; Pfaff et al., 2013), LaRocca et al., manuscript in preparation]. We compared the transcriptome of mouse KP cancer cells expressing either T6B-YFP (T6B^WT^-YFP) or a mutant version (T6B^MUT^-YFP) that cannot bind Argonaute proteins and is therefore inactive. As shown in **Figure 5f**, genes harboring peaks identified by HEAP were preferentially de-repressed upon disruption of the miRISC, further confirming the ability of the HEAP method to identify functional miRNA-mRNA interactions *in vivo*.

## DISCUSSION

We have demonstrated the ability of HEAP to identify miRNA-mRNA interactions sites in cells, developing embryos, normal adult tissues, and in primary autochthonous tumors. By mapping miRNA binding sites in mouse embryos lacking the miR-17∼92 cluster, we identified direct targets of the miRNAs encoded by this cluster, including a long noncoding RNA that had not been previously reported to be regulated by this cluster. The HEAP method also allowed us to identify miRNA targets in primary autochthonous cancers in mice and in their tissues of origin, uncovering marked differences in the spectrum of miRNA targets between cancers and normal tissues.

When compared to standard immunoprecipitation-based approaches, HEAP offers several advantages. First, the covalent nature of the interaction between the HaloTag and the HaloTag ligands simplifies the isolation of Ago2-miRNA-mRNA complexes and removes the intrinsic variability of immunoprecipitation-based approaches. This feature is illustrated by the highly reproducible identification of miRNA-binding sites in murine embryonic stem cells, in developing embryos, in murine tissues and in tumors. Second, the conditional Cre-lox-based nature of the Halo-Ago2 mouse strain enables the purification of Ago2-containing complexes and the identification of miRNA-mRNA interaction sites from a specific subset of cells, thus bypassing the need for microdissection and cell purification using cell surface markers. As proof of concept, we demonstrate this ability by mapping miRNA-mRNA interactions in three mouse models of human cancers driven by distinct combinations of oncogenes and tumor suppressor genes. We predict that the systematic application of HEAP will allow the construction of a detailed map of miRNA targets across tissues and cell types in mice.

We emphasize that the HEAP protocol can be easily modified to accommodate the many variations of the basic HITS-CLIP strategy, including those using ligation to generate chimeric reads between the mature miRNA and its target (CLASH, CLEAR-CLIP), and those designed to identify the cross-linking site at single base resolution (PAR-CLIP, iCLIP, eCLIP).

Although in this study we have focused exclusively on the identification of miRNA-mRNA interactions in cells and tissues, the conditional Halo-Ago2 mouse strain we have developed could prove useful for the biochemical characterization of Ago2-containing protein complexes *in vivo* and for imaging studies (**Supplementary Figure 14 and Supplementary Movie 1**). Notably, fluorescent HaloTag ligands have been successfully used recently for super-resolution imaging of Halo-tagged proteins (Grimm et al., 2015). When applied to cells and tissues expressing the Halo-Ago2 knock-in allele, this strategy could provide novel insights into the subcellular localization and dynamics of this important RNA binding protein under different conditions and in response to external and internal cues.

Despite these advantages, some limitations of the HEAP method should be considered when planning experiments. First, as is true for any tagged protein, the presence of the HaloTag may have functional consequences. The reduced viability of the Halo-Ago2 homozygous animal we have observed does indicate that Halo-Ago2 is not entirely functionally identical to Ago2. A possible explanation is that Halo-Ago2 fusion has reduced stability as compared to endogenous Ago2, since we have evidence that the Halo-Ago2 protein expressed from the endogenous Ago2 locus is incorporated into the miRISC complex, efficiently binds to miRNAs and target mRNAs, and retains slicing activity.

In conclusion, the HEAP method and the Cre-inducible Halo-Ago2 mouse strain described in this paper, combined with the growing array of strains expressing Cre in a temporally and spatially restricted fashion, will facilitate the generation of detailed maps of miRNA-mRNA interactions in vivo under physiologic and pathologic conditions.

## METHODS

### Cell culture

All cells were maintained in a humidified incubator at 37 °C, 5% CO_2_. mESCs were grown on irradiated MEFs in KnockOut DMEM (Gibco) supplied with 15% FBS (Gibco), leukemia inhibitory factor (Millipore, 10 U / mL), penicillin/streptomycin (Gibco, 50 U/ mL), GlutaMax (Gibco), non-essential amino acids (Sigma-Aldrich), nucleosides (Millipore) and 2-Mercaptoethanol (Bio-Rad, 100 µM). MEFs were cultured in DMEM (Gibco) containing 10% FBS (VWR), penicillin/streptomycin (100 U/mL) and L-glutamine. Mouse KP lung adenocarcinoma cells were cultured in Advanced DMEM/F12 (Gibco) containing 5% FBS, HEPES (Gibco, 10 mM), GlutaMax and penicillin/streptomycin (100 U/mL).

### Luciferase Assay

Ago2-null MEFs (obtained from Alexander Tarakhovsky, at the Rockefeller University) were transduced with the MSCV-PIG, MSCV-PIG-Halo, MSCV-PIG-Halo-Ago2 or MSCV-PIG-Ago2 retroviruses, to generate cell lines stably expressing HaloTag, the Halo-Ago2 fusion or Ago2. Dual-luciferase reporter assay system (Promega) was used to measure cleavage activity of Halo-Ago2 and Ago2. Luciferase reporter plasmids pIS0 (*luc+*, Firefly luciferase, (Yekta et al., 2004), Addgene: 12178) and pIS1 (*Rluc*, Renilla luciferase, Addgene: 12179) were co-transfected into MEFs, along with a pSico vector expressing an shRNA against the firefly luciferase or a control shRNA against CD8 (Ventura et al., 2004). The ratio between firefly and renilla luciferase activity was measured following manufacturer’s instructions at 48 hrs post-transfection.

### Mouse husbandry and generation of the conditional Halo-Ago2 knock-in mice

The targeting construct was generated by modifying the pKO-II vector through three steps of cloning. First, a fragment comprising a 2 kb 5’ homology arm, the 5’UTR of Ago2, the HaloTag cDNA, the TEV protease recognition sequence, the coding sequence of Ago2 Exon1 and a portion of the first intron was inserted into the pKO-II vector immediately upstream of the frt-PGK-NEO-frt cassette (between XhoI and AloI sites). Second, a 5 kb 3’ homology arm was cloned into the HindIII site downstream of the frt-PGK-NEO-frt casette. Lastly, a loxP-STOP-IRES-FLAG-loxP cassette was inserted into the AsiSI site between the TEV cleavage sequence and Ago2 coding sequence.

V6.5 mESCs (obtained from the Rudolf Jaenisch laboratory at Whitehead Institute and Massachusetts Institute of Technology) were electroporated with the linearized targeting construct and selected in mESC medium containing G418 for 7 days. Recombinant clones were identified by southern blot using probes designed against sequences outside the 5’ and 3’ homology regions. A validated clone was injected into C57BL/6 blastocyst to generate chimeric mice. Mice heterozygous for the targeted allele were crossed to the β-actin-Flpe mice (Rodriguez et al., 2000) to remove the frt-PGK-NEO-frt cassette, resulting in the generation of Ago2^Halo-LSL/+^ mice. The Ago2^Halo/+^ mice were obtained by crossing the Ago2^Halo-LSL/+^ mice to the CAGGS-Cre mice (Araki et al., 1997).

Mice carrying the knock-in alleles were genotyped using a three-primer PCR (p1, 5’-GCAACGCCACCATGTACTC-3’, final concentration 0.75 µM; p2, 5’-GAGGACGGAGACCCGTTG-3’, final concentration 1.0 µM, p3, 5’-AGCCGTTCCTGAATCCTGTT-3’, final concentration 0.5 µM), which amplifies a 240-bp band from the wild-type allele (p1-p2), a 1281-bp band from the Ago2^Halo-LSL^ allele and a 651-bp band from the Ago2^Halo^ allele (p2-p3). All studies and procedures were approved by the Memorial Sloan Kettering Cancer Center Institutional Animal Care and Use Committee.

### mESC mutagenesis

The Dicer1 knockout cells were generated from Ago2^Halo/+^ mESCs using CRISPR-Cas9. A pX333 vector ((Maddalo et al., 2014), Addgene: 64073) expressing Cas9 and a pair of guide RNAs designed to delete a portion of the RNase III 1 domain of Dicer1, was transiently transfected into Ago2^Halo/+^ mESCs. Single clones were isolated and genotyped by PCR.

Lefty2 mutant clones were generated from Ago2^Halo/+^ mESCs using CRISPR-Cas9-mediated homologous recombination. PX330 vectors ((Cong et al., 2013; Ran et al., 2013), Addgene: 42230) expressing Cas9 and guide RNAs targeting the predicted miR-291-3p binding site in the 3’UTR of Lefty2 were transiently transfected, together with single-stranded template DNA, into Ago2^Halo/+^ mESCs. Clones undergoing homologous recombination were enriched using the method developed by Flemr and Buhler (Flemr and Buhler, 2015). Clones homozygous for the desired mutations were identified by genotyping.

### Northern blotting

Total RNAs were isolated using TRIzol Reagent (Invitrogen) according to the manufacturer’s instructions. 20 µg total RNAs from samples were loaded into a 15% TBE-Urea polyacrylamide gel and transferred to a Hybond-N^+^ membrane (GE Healthcare). After UV crosslinking and blocking, a ^32^P-labeled DNA probe reverse complement to miR-293-3p was hybridized with the membrane at 37°C overnight. Next day, the membrane was washed and exposed to a film.

### Mass spectrometry proteomics

Five independent Dicer1 knockout and five wild-type mESC clones were used for the proteomic analysis. Frozen cells were lysed with lysis buffer containing 8 M Urea, 200 mM EPPS (pH8.5) and protease inhibitors. Total proteins were denatured and digested with LysC (20 µg/mL) and (10 µg/mL). Peptides were labeled with 0.2 mg TMT isobaric label reagents, pooled at a 1:1:1:1:1:1:1:1:1:1 ratio and subjected to mass spectrometry (MS). Raw MS data files were processed using MaxQuant (Cox and Mann, 2008). This resulted in a filtered matrix of protein abundance values for 8056 proteins. Then log2FC of abundance was calculated for each protein by summing values within five replicates of each condition, adding 1 to each sum, and then taking log2 of the ratio of the sums.

### MEF derivation

MEFs were derived from mouse E13.5 embryos following standard protocols. Ago2^Halo^ and Ago2^Halo-LSL^ MEFs were generated by intercrossing Ago2^Halo/+^ and Ago2^Halo-LSL/+^ mice, respectively. MEFs were immortalized with retrovirus expressing the SV40 large T antigen (Zhao et al., 2003) (Addgene:13970).

### Halo-Ago2 live cell imaging

Ago2-null MEFs transduced with retroviruses MSCV-PIG, MSCV-PIG-Halo or MSCV-PIG-Halo-Ago2 were treated with 5µM HaloTag TMR ligand (Promega) overnight and imaged with fluorescence microscope (ZEISS).

Ago2^+/+^, Ago2^Halo-LSL/Halo-LSL^ and Ago2^Halo/Halo^ MEFs were treated with 200 mM Janelia Fluor 646 HaloTag ligands or Janelia Fluor 549 HaloTag ligands (Promega) 1hr prior to experiment. Before imaging, medium containing Halo-ligand was replaced with warm medium without phenol red. Confocal imaging was performed using a ZEISS LSM880 microscopy with Airyscan super-resolution mode. A Plan-Apochromat 63x/1.4 Oil objective (Zeiss) was used. Time lapse images were acquired with a Zeiss alpha Plan-Apochromat 100X/1.46NA objective on an Axio Observer.Z1 in widefield using a Hamamatsu ORCA Flash4.0 v2 camera. Intervals between frames was either 0.5 or 1 seconds with 250ms exposures while at 37°C with 5% CO2 and 100% humidity incubation.

### Size exclusion chromatography

Cells were lysed with Sup6-150 buffer (150 mM NaCl, 10 mM TrisHCl, pH7.5, 2.5 mM MgCl_2_, 0.01% Triton X-100, protease inhibitor and phosphatase inhibitor). Lysates were fractionated using the Superose 6 10/300 GL prepacked column (GE Healthcare) coupled with the AKTA FPLC system as described in (La Rocca et al., 2015; Olejniczak et al., 2013). Eluted proteins were concentrated by trichloroacetic acid (TCA)/ acetone precipitation and analyzed by immunoblot.

### Isolation of Halo-Ago2/Tnrc6 complexes

Ago2^Halo-LSL/Halo-LSL^ and Ago2^Halo/Halo^ MEFs were lysed with HaloTag protein purification buffer (150 mM NaCl, 50 mM HEPES, pH7.5, 0.005% IGEPAL CA-630) and lysates were incubated with the HaloTag magnetic beads (Promega) for 90 min at room temperature on a rotator. After washes, proteins were released by TEV protease (Invitrogen) digestion at 30 °C for 1 hr. Eluted proteins were analyzed by immunoblot.

### Antibodies

Antibodies used were anti-Ago2 antibody (CST clone C34C6, 1:1000), anti-β-actin antibody (Sigma-Aldrich clone AC-74, 1:5000), anti-Tubulin antibody (Sigma-Aldrich clone DM1A, 1:5000), anti-HaloTag monoclonal antibody (Promega G9211, 1:1000), anti-Flag antibody (Sigma-Aldrich F7425, 1:1000), anti-Dicer1 (Bethy A301-936A, 1:1000) antibody and anti-TNRC6A (GW182) antibody (Bethyl A302-329, 1:1000).

### Recombinant adenovirus delivery

Recombinant adenoviruses used for inducing chromosomal rearrangement (Ad-BN, Ad-EA) (Cook et al., 2017; Maddalo et al., 2014) and Ad-Cre were purchased from ViraQuest.

For the generation of Bcan-Ntrk1-driven gliomas, a 1:1 mixture of Ad-BN and Ad-Cre, in total ∼3 × 10^9^ infectious particles, was administrated to Ago2^Halo-LSL/+^; p53^fl/fl^ mice (4∼6 weeks old), via stereotactic intracranial injection as described in Cook et al., 2017. Gliomas were harvested approximately 80 days after injection, when mice became symptomatic. For the generation of Eml4-Alk-driven lung adenocarcinomas, 10∼12-week-old Ago2^Halo-LSL/+^ mice were intratracheally infected with a 1:1 mixture of Ad-EA and Ad-Cre (in total ∼6 × 10^10^ infectious particles). To generate KRas^G12D^; p53-null lung tumors, 10∼12-week-old Ago2^Ha-lo-LSL/+^; KRas^LSL-G12D/+^; p53^fl/fl^ mice were intratracheally infected with AdCre (∼2.5 × 10^7^ PFU). Lung tumors were harvested approximately 3 months after infection.

### T6B peptide in KP lung adenocarcinoma cell lines

Mouse KP cells were derived from mouse KRas^G12D^, p53-null lung adenocarcinomas and transduced with retroviruses expressing the T6B^WT^-YFP or the T6B^MUT^-YFP, in which five tryptophan residues were mutated to alanine [(Hauptmann et al., 2015; Pfaff et al., 2013) and LaRocca et al., manuscript in preparation].

### RNA sequencing

Total RNAs from mESCs, lung adenocarcinomas and normal lung tissues were extracted using TRIzol Reagent and subjected to DNase (Qiagen) treatment followed by RNeasy column clean-up (Qiagen). After quantification and quality control, 500ng of total RNA underwent polyA selection and TruSeq library preparation using the TruSeq Stranded mRNA LT Kit according to the manufacturer’s instructions. Samples were barcoded and run on a HiSeq 2500 or a Hiseq 4000 in a 50bp/50bp paired end run. For mESCs, an average of 41 million paired reads were generated per sample. For lung and lung tumors, an average of 40 million paired reads were generated per sample.

Total RNAs of T6B-YFP-expressing KP cells were isolated using TRIzol Reagent and subjected to DNase treatment and isopropanol re-precipitation. After quantification and quality control, 1 ug of total RNA underwent ribosomal depletion and library preparation using the TruSeq Stranded Total RNA LT Kit. Samples were run on a HiSeq 4000 in a 50bp/50bp paired end run. On average, 34 million paired reads were generated per sample.

Reads were aligned to the standard mouse genome (mm10) using Hisat2 (v0.1.6-beta) (Kim et al., 2019) or STAR v2.5.3a (Dobin et al., 2013). RNA reads aligned were counted at each gene locus. Expressed genes were subjected to differential gene expression analysis by DESeq2 v1.20.0 (Love et al., 2014).

### Analysis of public datasets

RNA-seq data generated from E9.5 miR-17∼92 mutant embryos were obtained from the authors and are available in GEO ((Han et al., 2015), GSE63813). In this study, gene expression was profiled in triplicates in heart, mesoderm and all remaining tissues of wild-type (WT) embryos and embryos null for miR-17 and miR-20a (Δ17), null for miR-18a (Δ18), null for miR-19a and miR-19b-1 (Δ19), and null miR-92a-1 (Δ92), null for miR-17, miR-18a and miR-20a (Δ17,18), null for miR-17, miR-18a, miR-20a and miR-92a-1 (Δ17,18,92), and null for the entire cluster (KO). Embryos were of different genders. The data was aligned using HISAT v0.1.6-beta. In each tissue, differential gene expression analysis was performed using DESeq2 v1.6.3 using multi-factorial model “∼ d17 + d18 + d19 + d92 + gender”, where factor “d17” encoded for conditions that were Δ17, factor “d18” encoded for conditions that were Δ18, etc., and factor “gender” encoded for the genders of the embryos. This allowed us to estimate the log2FC of expression associated with each individual miRNA family in each tissue when accounting for contribution from other miRNA family and the gender.

The microarray dataset from CAD cell expressing miR-124 was obtained from ((Makeyev et al., 2007), GSE8498) using function getGEO() from GEOquery v2.50.5 (Davis and Meltzer, 2007). Differential expression analysis was run using functions lmFit() and eBayes() from limma v3.38.3 (Ritchie et al., 2015).

The iCLIP ((Bosson et al., 2014), GSE61348) and CLEAR-CLIP ((Moore et al., 2015), GSE73059) datasets were processed and aligned to the UCSC mm10 mouse genome using STAR v2.5.3a. Reads mapping to multiple loci or with more than 5 mismatches were discarded.

### HEAP and input control library preparation

mESCs were harvested and irradiated with UV at dose 400 mJ/cm^2^ in cold PBS on ice. Fresh tissues were harvested, homogenized and irradiated with UV for three times at dose 400 mJ/cm^2^. Cell or tissue pellets were snap frozen on dry-ice and stored at -80 °C.

Frozen pellets were thawed, lysed with mammalian lysis buffer (50 mM Tris-HCl, pH7.5, 150 mM NaCl, 1% Triton X-100 and 0.1% Na deoxycholate) containing protease inhibitor cocktail (Promega) and treated with RQ1 DNase (Promega). In order to get the “footprint” Halo-Ago2, lysates were treated with RNase A (1:50,000 diluted in TBS). ∼2% of the lysates were saved for input control library preparation. The remaining lysates were diluted with TBS (700 µL TBS per 300 µL lysates). For each sample, 300 µL Halolink resin (Promega) was used. The Halolink resin was equilibrated and incubated with the TBS-diluted lysates at room temperature for 1.5 hr. After incubation, the resin was washed extensively with a series of buffers: SDS elution buffer (50 mM Tris-HCl, pH7.5 and 0.1% SDS, one wash for 30 min at room temperature on a rotator), LiCl wash buffer (100 mM Tris-HCl, pH8.0, 500 mM LiCl, 1% IGEPAL CA-630 and 1% Na deoxycholate, three times), 1x PXL buffer (1x PBS with 0.1% SDS, 0.5% Na deoxycholate and 0.5% IGEPAL CA-630, two times), 5x PXL buffer (5x PBS with 0.1% SDS, 0.5% Na deoxycholate and 0.5% IGEPAL CA-630, two times) and PNK buffer (50 mM Tris-HCl, pH7.4, 10 mM MgCl_2_ and 0.5% IGEPAL CA-630, two times).

After dephosphorylation with calf intestinal alkaline phosphatase (Promega) and washes with buffer PNK-EGTA (50 mM Tris-HCl, pH7.4, 20 mM EGTA and 0.5% IGEPAL CA-630, two times) and PNK (two times), a 3’ RNA adaptor with a phosphate on its 5’ end (RL3) was ligated to the 3’ end of RNAs using T4 RNA ligase 1 (NEB) at 16 °C overnight. Next day, the resin was sequentially washed with the buffer 1x PXL (once), 5x PXL (once) and PNK (three times). RNAs on the resin were treated with T4 PNK (NEB) and washed with buffer PNK (three times), Wash/Eq (once) and PK (100 mM Tris-HCl, pH7.5, 50 mM NaCl and 10 mM EDTA, once). To release RNAs from the resin, proteins were digested with 4 mg/mL proteinase K (Roche) in PK buffer and further inactivated by 7 M urea dissolved in PK buffer. Free RNAs were extracted using phenol/chloroform and precipitated with ethanol/ isopropanol at -20 °C overnight. Next day, RNAs were pelleted, washed with 70% cold ethanol and resuspended in DEPC-treated H_2_O. A 5’ RNA adaptor (RL5) with six degenerate nucleotides and a common ‘G’ on its 3’ end (RL5-NNNNNNG, RL5D-6N) was ligated to the purified RNAs using T4 RNA ligase 1 at 16 °C for 5 hrs. Then, the RNAs were treated with RQ1 DNase to remove residual DNAs and purified by phenol/chloroform extraction and ethanol/isopropanol precipitation.

Purified RNAs were reverse transcribed using the DP3 primer (final concentration: 0.5 µM) and Superscript III reverse transcriptase (Invitrogen). The resulting cDNAs were amplified with primers DP3 and DP5 (final concentrations: 0.5 µM) to the optimal amplification point. The optimal amplification cycle (defined as the cycle before the PCR reaction reaching a plateau) was preliminarily determined by a diagnostic PCR visualized on gel or by a real-time PCR with SYBR green. PCR products of miRNAs (HEAP miRNA library, expected size: 65 bp) and targets (HEAP mRNA library, expected size range: 75∼200 bp) were resolved on a 15% TBE-Urea polyacrylamide gel and extracted separately (**Supplementary Figure 2a**). To construct library for high-throughput sequencing, DNA primers DP3-barcodeX and DSFP5 (final concentrations: 0.5 µM) containing Illumina adaptors, sequencing primer binding sites and Illumina TruSeq indexes for multiplexing were introduced to the HEAP miRNA and mRNA libraries by PCR.

To prepare input control library, RNAs in the lysates saved before the Halolink resin pulldown were dephosphorylated with calf intestinal alkaline phosphatase and phosphorylated using with T4 PNK. RNAs were then cleaned up using the MyONE Silane beads (ThermoFisher Scientific) as described in (Van Nostrand et al., 2016). Then, the 3’ RNA adaptor (RL3) was ligated to the purified RNAs at 16 °C overnight. Next day, the ligated RNAs were purified using the MyONE Silane beads. Similar to the preparation of HEAP libraries, the RNAs were ligated to the 5’ RNA adaptor (RL5D-6N) at 16 °C for 5 hrs, treated with RQ1 DNase, purified, reverse transcribed to cDNAs and amplified by PCR using primers DP3 and DP5. PCR products ranging from 75 to 200 bp were resolved on a 15% TBE-Urea polyacrylamide gel and used as input libraries.

### High-throughput sequencing of the HEAP and input control libraries

HEAP mRNA and miRNA libraries, along with the matched input control libraries, were submitted to the Integrated Genomics Operation Core at Memorial Sloan Kettering Cancer Center for high-throughput sequencing. After quantification and quality control, libraries were pooled and run on a HiSeq 2500 in Rapid mode in a 100 bp or 125 bp single end run, using the HiSeq Rapid SBS Kit v2 (Illumina).

### HEAP library read processing, alignment and deduplication

#### Step 1: Barcode removal

The 6 nt degenerate barcodes and the last nucleotide ‘G’ coming from the 5’ adaptor RL5D-6N (in total 7 nt) were removed from the beginning of reads and appended to the original read names, which later were used to distinguish duplicated reads produced at PCR amplification steps.

#### Step 2: Adaptor removal and read quality control

The 3’ adaptor (5’-GTGTCAGTCACTTCCAGCGGGATCGGAAGAG-CACACGTCTGAACTCCAGTCAC-3’) and bases with Phred quality score lower than 20 were trimmed from reads using cutadapt v1.15 or v1.17. After trimming, reads shorter than 18 nt were discarded.

#### Step 3: Alignment

Processed reads were aligned to the UCSC mm10 mouse genome using STAR v2.5.3a. Reads mapping to multiple loci or with more than 5 mismatches were discarded.

#### Step 4: PCR duplicate removal

Reads mapped to the same locus with identical barcodes were considered PCR duplicates and therefore collapsed. This was achieved by storing aligned reads using chromosome names, strand information, positions of the first bases and the 7 nt barcodes as keywords. Representative reads of these unique events were written into a BAM file, which was used for peak calling.

### Peak calling

Peak calling and preprocessing was done using our package CLI-Panalyze (https://bitbucket.org/leslielab/clipanalyze). The function findPeaks() was used to run multiple steps of analysis. First, the combined signal from uniquely aligned and PCR-duplicate-corrected reads from multiple replicates was convolved with the second derivative of a Gaussian filter. Zero-crossings of the convolved signal corresponded to edges of putative peaks. Second, read counting was run in putative peaks and in GENCODE-annotated gene exons with putative peaks subtracted, for both HEAP replicates and input control replicates. Library sizes for both HEAP and input control replicates were estimated using the read counts in exons outside of putative peaks. Third, using these library size estimates, differential read count analysis was performed between HEAP and input control read counts in putative peaks using DESeq2, and FDR-corrected *p*-values (adjusted *p-*values) were assigned to each peak. Peaks of size > 20nt and read count log2FC > 0 in HEAP vs. control were selected for downstream analysis. Peaks were annotated as overlapping with 3’UTR, 5’UTR, exons, introns, intergenic regions, lncRNA, in that order, using GENCODE (vM17) annotation. Peaks overlapping with genes of types “lincRNA”, “antisense”, “processed_transcript”, according to GENCODE, were annotated as lncRNA peaks.

For mESCs, peak calling was run using three HEAP libraries against two input control libraries with the following parameters in findPeaks(): count.threshold = 10, extend.slice = 10, bandwidth = 80, extend.peaks. in.genes = 150. The same parameters were used when calling peaks for each individual replicate to assess reproducibility (**Supplementary Figure 2**) and for the Lefty2 wild-type and mutant libraries. For iCLIP, peak calling was run using a single iCLIP library (TT-FHAGO2) against a single control library (TT-AGO2) with the following parameters: count.threshold = 5, extend.slice = 50, bandwidth = 60, extend. peaks.in.genes = 150. For comparison with iCLIP, peak calling with the same parameters was run for each single HEAP library of comparable size against a single input control library (**Figure 2f, Supplementary Figure 5b-c**).

For embryos, peak calling was run using HEAP in one wildtype (miR-17∼92-WT), two heterozygous (miR-17∼92-HET) and one homozygous knockout (miR-17∼92-KO) embryo against the four matching input control libraries using the following parameters: count.threshold = 5, extend.slice = 10, bandwidth = 80, extend.peaks.in.genes = 150. Then differential HEAP read count analysis was performed using DESeq2 v1.22.1 in miR-17∼92-KO against miR-17∼92-WT and miR-17∼92-HET libraries to determine miR-17∼92-dependent peaks (plotted in **Supplementary Figure 8a**).

For cortices of P13 mice, peak calling was run using the two HEAP libraries against the two matching input control libraries using the same parameters as for embryos. The same parameters were used for peak calling using CLEAR-CLIP in 12 replicates vs. the input control libraries generated for HEAP. Differential HEAP read count analysis in HEAP vs. input was performed using DESeq2 v1.20.0.

For gliomas and cortices in adult mice, three HEAP libraries from each context were generated. Before peak calling, size factors Y of the six HEAP libraries were estimated using the byte sizes of corresponding BAM files. Then, BAM files for two glioma replicates and three cortex replicates were downsampled to similar sizes to the smallest glioma replicate using samtools v1.3.1 (Li et al., 2009) with scaling factors X = 1/Y. Peak calling was run using the six scaled HEAP libraries against the six matching input control libraries, using the same parameters as for embryos, to identify the set of putative peaks. As usual, only peaks of size > 20nt and with log2FC > 0 in HEAP vs. control were used in downstream analysis. Furthermore, only peaks with average normalized read count > 10 in the three glioma replicates or in the three cortex replicates were selected. To identify significant peaks in gliomas, DESeq2 v1.20.0 for read counts in these selected peaks was run using the three glioma replicates against the three matching input control replicates. To identify significant peaks in cortices, DESeq2 for read counts in these selected peaks was run using the three cortex replicates against the three matching input control replicates. Differential HEAP read counts analysis between gliomas and cortices was run in peaks with adjusted *p-*value < 0.05 (in HEAP vs. control).

For lung tumors, peak calling was run using two HEAP libraries generated from normal lungs, two HEAP libraries from KP tumors and three HEAP libraries from EA tumors against seven matching input control libraries, using the same parameters as for embryos. Peaks of size > 20nt and with log2FC > 0 in HEAP vs. control were used in downstream analysis. Furthermore, only peaks with average normalized read count > 10 in the two normal lung replicates, in the two KP tumor replicates or in the three EA tumor replicates were selected. To identify significant peaks in each tumor type, DESeq2 v1.20.0 for read counts in these selected peaks was run using the tumor replicates against their matching input control replicates. To identify significant peaks in normal lungs, DESeq2 for read counts in the selected peaks was run using the two normal lung replicates against the two matching input control replicates. To compare peak intensities between KP and EA tumors, DESeq2 for read counts in peaks with adjusted *p-*value < 0.05 (in HEAP vs. input) was run using the three EA tumor replicates against the two KP tumor replicates. Since peak intensities in EA and KP highly correlate with each other, the five tumor replicates were grouped and used for downstream analysis. To compare peak signals between tumors and normal lungs, differential HEAP read count analysis was perform in peaks with adjusted *p*-value < 0.05 (in HEAP vs. input) between the five tumor replicates and the two normal lung replicates.

### miRNA abundance estimates

Reads in the HEAP miRNA libraries were processed and filtered following step 1 and step 2 described in the “HEAP library read processing, alignment and deduplication” section. Processed small RNA reads were aligned to a miRNA genome index built from 1,915 murine pre-miRNA sequences from miRbase version 21 (ftp://mirbase.org/pub/mirbase/21/) using Bowtie v2.3.4 (Langmead and Salzberg, 2012), and these reads were considered true miRNA counts if they fell within ± 4 bps at each of the 5’ and 3’ end of the annotated mature miRNAs. PCR duplicates were removed as described in step 4 in the “HEAP library read processing, alignment and deduplication” section.

miRNA seed family data were downloaded from the TargetScan website at http://www.targetscan.org/mmu_71/mmu_71_data_download/miR_Family_Info.txt.zip. For miRNA family level analysis, read counts mapping to members of the same miRNA family were summed up.

### mRNA abundance estimates using input control libraries

Input control libraries generated from gliomas and cortices were used to estimate mRNA abundance. Reads were counted at each gene locus using featureCounts v1.6.3 (Liao et al., 2014) with GENCODE (vM22) primary annotation. Differential gene expression analysis was performed using DEseq2 v1.20.0.

### Motif discovery

#### 1. Unbiased motif enrichment analysis

Frequencies of all k-nucleotide-long sequences (k-mers, k = 7) were calculated for sequences in selected peaks (Freq_selected_) and background sequences (Freq_bg_). The enrichment score for these 7-mers was calculated as log2FC = log2 ((Freq_selected_ + c) / (Freq_bg_ + c)), where c was a small corrective value that dependeds on k, the number and size of peaks. K-mers with the highest log2FCs were then reported. This analysis was performed using functions calculateKmerBackground() and findKmerEnrich() in CLIPanalyze. For mESCs, peaks mapping to 3’UTR were selected and background sequences were defined as sequences of 3’UTRs outside of peaks. For brain and lung cancers, peaks differentially present in tumors and their tissues of origin (adjusted *p*-value < 0.1, absolute log2FC (tumor vs. normal) > 0.5) were selected and compared against background sequences, defined as exon sequences of genes, where peaks were identified.

#### 2. Enrichment score calculation for miRNA seed matches

log2 enrichment score of miRNA seed matches in **Figure 2b** was calculated as log2 (Freq_3’UTR_ / Freq_bg_). Freq_3’UTR_ was frequencies of 8mer seed matches for miRNA seed families in 3’UTR peaks, while Freq_bg_ was frequencies of these seed matches calculated in background sequences. Background sequences were defined as 3’UTR sequences outside of peaks.

#### 3. HOMER *de novo* motif discovery

In mESC libraries, for the top 50 7-mers found by unbiased motif enrichment analysis, positions of their exact occurrences in 3’UTR peaks were found. Sequences of a 15-bp region around these occurrences were extracted and subjected to HOMER *de novo* motif discovery (Heinz et al., 2010), using 15-bp windows shifted 100 bp and 200 bp on both sides of the 7-mers (and excluding those overlapping with any of the 3’UTR peaks) as background sequences. Similarly, for glioma and cortex libraries, the top 50 7-mers found in each context were mapped to corresponding peak sets and subjected to HOMER *de novo* motif discovery. For normal lung and lung tumor libraries, the top 70 7-mers from each context were used.

### IDR analysis

IDR analysis was run using the python package at https://github.com/nboley/idr (Li et al., 2011). All putative peaks (size > 20nt, log2FC > 0 for HEAP vs. control) were provided via parameter “--peak-list”. Peaks called for individual replicates were scored using log2FC in HEAP vs. control and provided via parameter “--samples”, separately for each pair of replicates. Peaks at IDR < 0.05 were considered reproducible.

### HEAP coverage analysis

bigWig files for visualization of HEAP and input control libraries at 1bp resolution were produced in the following way. First, deepTools bamCoverage v3.1.3 (Ramirez et al., 2016) with parameter “-bs 1 --scale-Factor X” was used to produce bedGraph files. Here, size factors Y were estimated using DESeq2 applied to read counts in exons outside of peaks in all HEAP and input control libraries in a particular experimental model (mESCs, embryos, etc.) and then reciprocals X = 1 / Y were used as BAM coverage scaling factors. Only bedGraph signal in the standard chromosomes was selected. Then “bedtools sort” (using bedtools v2.23.0 (Quinlan and Hall, 2010)) and bedGraphToBigWig v4 (Kent et al., 2010) were used to produce bigWig files.

To measure HEAP coverage of various peak sets in embryo libraries, peaks were first assigned to miRNA seed families by searching for the corresponding 7mer and 8mer seed matches in peak sequences and then grouped based on their genomic locations. Score matrices of 800-bp windows surrounding these peaks were generated from size-factor-corrected bigWigs using the ScoreMatrixList() function from the genomation package v1.14.0 (Akalin et al., 2015). Histograms of average score were produced using the function plotMeta(). Heatmaps were generated using the multiHeatMatrix() function and extreme values were removed before plotting using the winsorize parameter with values c(0,98).

## Supporting information

Computational pipeline

Supplementary protocol

## ACKNOWLEDGMENTS

We acknowledge the use of the Integrated Genomics Operation Core, funded by the NCI Cancer Center Support Grant (CCSG, P30 CA08748), Cycle for Survival, and the Marie-Josée and Henry R. Kravis Center for Molecular Oncology. This work was funded by grants from the NIH (NCI R01CA149707), The Starr Cancer Consortium, and the Geoffrey Beene Cancer Research Foundation to AV. XL was funded by the Geoffrey Beene Graduate Student Fellowship (2015∼2016). YP was supported by the 2019 AACR-Bristol-Myers Squibb Immunooncology Research Fellowship, Grant Number 19-40-15-PRIT. CPC was supported by the NCI F31 training grant (2013∼2015), Grant Number 1F31CA168356-01A1.

We thank Gregory Hannon for suggesting the use of the Halo-tag, Joana de Campos Vidigal for assistance with the design of the Halo-Ago2 targeting construct and the gene targeting experiments, and members of the Ventura, Leslie, and Benezra laboratories for discussion and suggestions.

## DATA SHARING

RNAseq and HEAP datasets are available for download from GEO with accession number GSE139349. Reagents and the mouse strains described are available upon request. Bigwig tracks can for several HEAP experiments described in this paper can be accessed at http://genome.ucsc.edu/s/yuri.pritykin/mm10%20HEAP%20ES%2C%20brain%2C%20lung%20updated

