## Supplementary material for "High-resolution *in vivo* identification of miRNA targets by Halo-Enhanced Ago2 Pulldown": Computational pipeline

### HEAP and input library processing pipeline

#### Step 1. Barcode removal

Python script: barcode.py

```
#!/usr/bin/env python2
import multiprocessing
import functools
from Bio import Seq, SeqIO
import gzip
import csv
import os
import glob

def attach_barcode( file_name, barcode_len ):
    input_file = "%s.fastq.gz" % file_name
    input_handle = gzip.open( input_file, "rb" )
    output_file = "%s_barcode.fastq.gz" % file_name
    output_handle = gzip.open( output_file, "wb" )

    for record in SeqIO.parse( input_handle, "fastq" ):
        record.description = ""
        sequence = str( record.seq )
        tag = sequence[ :barcode_len ]
        record.id = record.id + ":" + tag
        record = record[ barcode_len: ]
        SeqIO.write( record, output_handle, "fastq" )

    print "%s Complete!\n" % file_name
    input_handle.close()

file_names = glob.glob("*fastq.gz")
file_names = [ x[:-9] for x in file_names ]

max_num_process = len( file_names )
pool = multiprocessing.Pool( max_num_process )
test = pool.map( functools.partial( attach_barcode, barcode_len = 7 ),
file_names )
```

Usage: Barcode length (“barcode_len”) is 7. The 6-nt degenerate barcode and the “G” introduced by RL5D-6N is to be removed.

#### Step 2. Adaptor removal and quality trimming

Script:

```
cutadapt -a GTGTCAGTCACTTCCAGCGGGATCGGAAGAGCACACGTCTGAACTCCAGTCAC -m 18 -q
20 -o filename.trimmed.fastq.gz filename.fastq.gz
```

#### Step 3. Alignment (for HEAP mRNA and input control libraries only)

Script:

```
STAR \
      --genomeLoad NoSharedMemory \
      --genomeDir mm10_star/ \
      --readFilesIn filename.trimmed.fastq.gz \
      --runThreadN 2 \
      --alignIntronMin 70 \
      --alignIntronMax 100000 \
      --outSAMtype BAM SortedByCoordinate \
      --outFilterMultimapNmax 1 \
      --outFilterMultimapScoreRange 0 \
      --outFilterMismatchNmax 5 \
      --outFileNamePrefix filename/ \
      --outStd Log \
      --readFilesCommand zcat \
      --outReadsUnmapped Fastx
```

Usage: “mm10_star” genomic index was preliminarily generated using command “STAR --runMode genomeGenerate”

#### Step 4. PCR duplicate removal

Perl script: remove_duplicate.pl

```
#!/usr/bin/env perl

# Collapse CLIP reads mapped to the same position & with the same barcode
use warnings;
use 5.014;

# Calculate the range that a read span
sub reallen {
    my $reallen = 0;
        while( $_[0] =~ m/(\d+)([MIDNS])/g ){
        my $val = $1;
        my $code = $2;
        given($code) {
            when ('M') { $reallen = $reallen + $val;}
            when ('I') { $reallen = $reallen + 0;}
            when ('D') { $reallen = $reallen + $val;}
            when ('N') { $reallen = $reallen + $val;}
            when ('S') { $reallen = $reallen + 0;}
        }
        }
        $reallen;
}

$\ = "\n";
# open FILE, $ARGV[0] or print "File $ARGV[0] doesn't exist.";
# Fields I need
my %reads = ();
my $tag = '';
```

```
my $chr = '';
my $strand = '';
my $pos = 0;
my $nh = '';
my %flag = ();
my $key = '';
while(<>){
    chomp;
        # Do nothing with the header lines
        if ( substr($_,0,1) eq "@"){
                print;next;
        }
        # Now this line is a real read
    my @fields = split "\t";
        # Skip unmapped reads
        next if($fields[2] eq '*');
        # Fields to keep:
    $tag = (split( ':', $fields[0] ))[-1];
        $chr = $fields[2];
        $strand = ( ($fields[1] & 0x10) == 0 )?"+":"-";
        # Position that the first nt of a read is mapped to
        # Positive strand: original pos
        # Reverse: go forward a bit
        $pos = ( ($fields[1] & 0x10) == 0 )?($fields[3]):( $fields[3] +
reallen($fields[5]) - 1 );
        $flag{ ( split ":", $fields[$_] )[0] } = ( split ":", $fields[$_]
)[-1] foreach ( 11 .. $#fields );
        $nh = $flag{NH};
        # die "Parsing the wrong field!\n" if ( ( split ":", $fields[13]
)[0] ne "X0" );
        $key = join ":", $chr, $strand, $pos, $tag;
        # $pos = ( ($fields[1] & 0x10) == 0 )?($fields[3]):( $fields[3] +
reallen($fields[5]) - 1);
        # Multi-hit reads have lower priority
        $reads{$key} = join "\t", @fields if( $nh == 1 or !exists(
$reads{$key} ) )
}
foreach( keys %reads ){
        print $reads{$_};
}
# close FILE;
```

#### Step 5. HEAP miRNA library alignment

Script:

```
bowtie2 --no-unal -p 7 -x mouse_hairpin/mouse_hairpin -U
filename.trimmed.fastq.gz | samtools view -bS - | samtools sort -o
filename.bam
```

Usage: “mouse_hairpin” miRNA genome index was built from 1,915 murine pre-miRNA sequences from miRbase version 21.

#### Step 6. miRNA abundance measurement

RScript: mature_mir_count.R

```
require( GenomicAlignments )
require( stringr )
require( parallel )

mir <- readRDS( "hairpin_info.rds" )
mir.end <- resize( resize( mir, 1, fix = 'end' ) , 9, fix = 'center' )
mir.end <- split( mir.end, mir.end$mature )

fl <- Sys.glob( "*.bam" )
aln <- lapply( fl, readGAlignments, param = ScanBamParam( tag = "NM" ) )


mir.counts <- mclapply( aln, function( aln, mir.end ){
    aln <- aln[ mcols( aln )$NM <= 1]
    gr <- resize( granges( aln ), 1, fix = "end" )

    counts <- countOverlaps(  mir.end, gr )
    return( counts )
} , mir.end, mc.cores = 7 )
mir.counts <- do.call( 'cbind', mir.counts )
colnames( mir.counts ) <- substr( basename(fl), 1, nchar(basename(fl)) - 4 )
write.csv( mir.counts, file = "mature_mir_counts_all.csv" )
```

Usage:
1. Call this Rscript from command line.
2. “hairpin_info.rds” is a GenomicRanges object containing coordinates of mature miRNAs relative to their pre-miRNAs.

#### Step 7. Peak calling

RScript

```
peak.data <-
    findPeaks(bamfiles,
              names(filenames),
              condition = c(1, 0, …),
              annot.order = c("utr3", "utr3*", "utr5", "utr5*",
                              "exon", "intron"),
              exclude.mirna.peaks = TRUE,
              genome = "mm10",
              bandwidth = bandwidth,
              count.threshold = count.threshold,
              extend.slice = extend.slice,
              count.exons.only = FALSE,
              extend.peaks.in.genes = extend.peaks.in.genes,
              transform.chr.names = TRUE,
              nthreads = 14)
```

Usage:

1. “bamfiles” is a vector containing all BAM files, including HEAP mRNA library and input library.
2. “filenames” is a vector of sample names matched to BAM files.
3. The “condition” parameter specifies whether the BAM file is a HEAP mRNA library (1) or an input library (0).
4. “bandwith”, “extend.slice”, “count.threshold” and “extend.peaks.in.genes” parameters used in each analysis were described in the Methods session.
