## Supplementary protocol for "High-resolution *in vivo* identification of miRNA targets by Halo-Enhanced Ago2 Pulldown"

### Halo-enhanced Ago2 pulldown (HEAP)

#### HEAP Library preparation

##### Step 1. Cell preparation and UV crosslinking

1. Harvest and irradiate cells.

For mESCs

a) Dissociate mESCs from culture dishes and wash with cold PBS.
b) Resuspend cells in cold PBS in a 10 $cm^2$ dish and place on ice.
c) Crosslink cells by placing the dish on ice in a Spectroline UV crosslinker. Irradiate cells at dose 400 $mJ/cm^2$

For fresh tissues or tumors

a) Euthanize mice following the standard protocol.
b) Resect tissues of interest from mice and place in cold PBS on ice.
c) Homogenize tissues with a scalpel.
d) Crosslink tissues on ice at dose 400 $mJ/cm^2$ for three times.

2. Collect cells or tissues by centrifugation at 4 °C, 1300 xg for 10 min.
3. Remove supernatant. Snap-freeze the pellet on dry ice and store at -80 °C until use.

##### Step 2. Lysates preparation and RNase digestion.

1. Resuspend pellets in 3 volumes of Mammalian Lysis Buffer containing protease inhibitor cocktail (Promega G6521).
2. Pipette/vortex to mix. Incubate on ice for 15 min.
3. Add 25 $\mu$L RQ1 DNase (Promega M6101) per 300 $\mu$L lysate. Incubate in a thermomixer at 37 °C, 1,000 rpm for 5 min.
4. Per 300 $\mu$l lysates, add 2.5 $\mu$L RQ1 DNase and 10 $\mu$L RNase A (Affymetrix, 1:50,000 diluted in TBS). Incubate in a thermomixer at 37 °C, 1,000 rpm for 5 min.
5. Pass lysates through a 26-gauge needle to reduce viscosity.
6. Clear lysates by centrifugation at 4 °C, 14000 rpm for 10 min. Transfer lysates to a new tube and place on ice.

##### Step 3. Halolink Resin equilibration

1. Homogenize the Halolink resin (Promega G1914) slurry by inversion and dispense into a 15 mL conical tube (300 $\mu$L slurry per sample).
2. Wash resin with 4 volumes of Wash/Eq buffer.
3. Collect resin by centrifugation at 800 xg for 2 min.
4. Repeat step 2-3 twice.

5. Leave the resin in Wash/Eq buffer in a 1.5 mL eppendorf tube until lysates are ready.

#### Step 4. Halo-Ago2 pulldown and washing

1. Save ~2% lysate for **input control library** preparation. Store at -80 °C until use.
2. Dilute rest of the lysates with TBS at a 3:7 dilution ratio (700 $\mu$L TBS added into 300 $\mu$L lysates).
3. Remove Wash/Eq buffer from the resin and add diluted lysates.
4. Incubate lysates with resin on a tube rotator at room temperature for a total of 1.5 hr. If the volume of diluted lysates is greater than 1 mL, collect the resin by centrifugation (at 800 xg for 2 min) and reload the resin with the same sample.
5. Wash resin with 100 $\mu$L SDS elution buffer. Rotate at room temperature for 30 min. Collect resin by centrifugation at 800 xg for 2 min.
6. Wash resin 3x with 1 mL LiCl wash buffer, 2x with 1 mL PXL (1x) buffer, 2x with 1 mL PXL (5x) buffer and 2x with 1 mL PNK buffer. Collect resin by centrifugation in between.

#### Step 5. Dephosphorylation and 3' RNA adaptor ligation

1. Prepare the following reaction mix:

| Ref | Components | Volume ($\mu$L) |
|---|---|---|
| Promega M183A | 10X Alkaline phosphatase buffer | 8 |
| Promega M182A | Calf Intestinal Alkaline Phosphatase | 3 |
| Promega N251B | rRNasin | 2 |
| | Water | 67 |

2. Add 80 $\mu$L mixture to the resin. Incubate in a thermomixer at 37 °C for 20 min. Mix at 1,000 rpm for 15 s every 2 min.
3. Wash resin 2x with 1 mL PNK-EGTA buffer and 2x with 1 mL PNK buffer.

#### Step 6. 3' RNA linker ligation

1. Prepare the following reaction mix:

| Ref | Components | Volume ($\mu$L) |
|---|---|---|
| NEB B0216S | 10X T4 RNA ligase buffer | 8 |
| | BSA (0.2 $\mu$g/$\mu$L) | 8 |
| NEB P0756S | ATP (10 mM) | 8 |
| NEB M0204S | T4 RNA ligase 1 | 3 |
| Promega N251B | rRNasin | 2 |
| | RL3 (20 $\mu$M) | 5 |
| | Water | 46 |

2. Add 80 μL mixture to the resin. Incubate in a thermomixer at 16 °C overnight. Mix at 1,000 rpm for 15 s every 2 min.

#### Step 7. PNK treatment

1. Next day, wash resin 1x with 1 mL PXL (1x) buffer, 1x with 1 mL PXL (5x) buffer and 3x with 1 mL PNK buffer.
2. Prepare the following reaction mix:

| Ref | Components | Volume (μL) |
|---|---|---|
| NEB B0201S | 10X PNK buffer | 8 |
| NEB P0756S | ATP (10 mM) | 1 |
| NEB M0201L | T4 PNK | 4 |
| Promega N251B | rRNasin | 2 |
| | Water | 65 |

3. Add 80 μL mixture to the resin. Incubate in a thermomixer at 37 °C for 20 min. Mix at 1,000 rpm for 15 s every 2 min.
4. Wash resin 3x with 1 mL PNK buffer and 1x with 1 mL Wash/Eq buffer.

#### Step 8. RNA isolation from resin

1. Dissolve proteinase K (Roche 3115836001) in PK buffer to a final concentration of 4 mg/mL. Pre-warm this solution in a thermomixer at 37 °C, 1,000 rpm for 20 min.
2. Wash the resin with 1 mL PK buffer.
3. Add 200 μL pre-warmed proteinase K solution. Incubate in a thermomixer at 37 °C, 1,000 rpm for 20 min.
4. Add 200 μL PK-Urea solution. Incubate in a thermomixer at 37 °C, 1,000 rpm for 20 min.
5. Add 400 μL Phenol (pH4.3, Sigma P4682) and 130 μL Chloroform (Sigma-Aldrich: 25668). Incubate in a thermomixer at 37 °C, 1,000 rpm for 20 min.
6. Spin at 14,000 rpm at room temperature for 5 min.
7. Transfer the aqueous phase to a siliconized tube.
8. Add 50 μL 3 M Sodium Acetate (pH 5.5, Ambion AM9740) and 0.75 μL Glycogen (5 mg/mL, Ambion AM9510). Mix thoroughly.
9. Add 1 mL cold Ethanol/Isopropanol (1:1 by volume). Precipitate overnight at -20 °C.

#### Step 9. 5' RNA linker ligation

1. Precipitate RNA by centrifugation at 4 °C, 14,000 rpm for 20 min.
2. Wash the pellet once with 1 mL cold 70% Ethanol. Spin at 4 °C, 14,000 rpm for 10 min.
3. Remove supernatant and air-dry the pellet.

4. Resuspend RNA pellet in 5.9 μL RT-PCR grade water.
5. Prepare the following reaction mix:

| Ref | Components | Volume (μL) |
|---|---|---|
| NEB B0216S | 10X T4 RNA ligase buffer | 1 |
| | BSA (0.2 μg/μL) | 1 |
| NEB P0756S | ATP (10 mM) | 1 |
| NEB M0204S | T4 RNA ligase 1 | 0.1 |
| | RL5D-6N (20 μM) | 1 |

6. Add 4.1 μL mix to dissolved RNA. Incubate at 16 °C for 5 hrs.

#### Step 10. DNase treatment

1. Prepare the following reaction mix:

| Ref | Components | Volume (μL) |
|---|---|---|
| Promega M198A | RQ1 DNase 10X Reaction Buffer | 11 |
| Promega N251B | rRNasin | 5 |
| Promega M610A | RQ1 DNase | 5 |
| | Water | 79 |

2. Add 100 μL mix to each ligation reaction. Incubate in a thermomixer at 37 °C, 1,000 rpm for 20 min.
3. Add 300 μL $H_2O$, 50 μL 3 M Sodium Acetate (pH 5.5) and 0.75 μL Glycogen (5 mg/mL). Mix thoroughly.
4. Add 1 mL cold Ethanol/Isopropanol (1:1 by volume). Precipitate overnight at -20 °C.

#### Step 11. Reverse-transcription

1. Precipitate RNA as described in Step 9 (1-3). Resuspend RNA in 10 μL RT-PCR grade water.
2. Prepare the following mix. (RT-: control reaction without reverse transcriptase)

| Ref | Components | RT+ | RT- |
|---|---|---|---|
| | DP3 (10 μM) | 1 | 1 |
| Invitrogen 18427-013 | 10 mM dNTP mix | 1 | 1 |
| | Water | 3 | 9 |

3. Add 5 μL RT+ mixture to 8 μL RNA and 11 μL RT- control mixture to 2 μL RNA.
4. Incubate at 65 °C for 5 min.
5. Prepare the following reaction mix:

| Ref | Components | RT+ | RT- |
|---|---|---|---|
| Invitrogen 18080044 | 5X First-Strand buffer | 4 | 4 |
| Invitrogen 18080044 | 0.1M DTT | 1 | 1 |
| Invitrogen 10777019 | RNaseOUT (40U/$\mu$l) | 1 | 1 |
| Invitrogen 18080044 | SuperScript III Reverse transcriptase (200U/$\mu$l) | 1 | 0 |
| | Water | 0 | 1 |

7. Add 7 $\mu$L enzyme-buffer mixture to corresponding RNA mixtures.
8. Incubate the reaction mixtures in a thermocycler using program:
   50 °C 45 min
   55 °C 15 min
   90 °C 5 min
   4 °C ∞

#### Step 12. First PCR to determine optimal amplification cycles

OPTION 1

1. Prepare the following PCR reaction mix:

| Ref | Components | Volume ($\mu$L) |
|---|---|---|
| Invitrogen 12344040 | Accuprime Pfx SuperMix | 13.5 |
| | DP5 (20 $\mu$M) | 0.375 |
| | DP3 (20 $\mu$M) | 0.375 |
| | cDNA* | 1 |

* Prepare separate mixtures for control samples (RT- and water).

2. Amplify the cDNAs on a thermocycler using the following program with several different cycles. For example, one can start with 18, 22 and 26 cycles.

| Temperature | Time | |
|---|---|---|
| 95 °C | 2 min | |
| 95 °C | 20 s | X cycles |
| 58 °C | 30 s | |
| 68 °C | 20 s | |
| 68 °C | 5 min | |
| 4 °C | ∞ | |

3. Mix the PCR product with 2x TBE-Urea sample buffer (Invitrogen LC6876).
4. Load the same sample amplified by different number of cycles next to each other. Run PCR products on a 15 % TBE-Urea polyacrylamide gel (Invitrogen EC6885), along with the 25-bp DNA step ladder (Promega G4511), following standard protocol.

5. Stain the gel in 1x SYBR gold nucleic acid gel stain (Invitrogen S11494) in 1 X TBE for 10 min.
6. Visualize the PCR product under UV. Expected size for miRNAs is ~65 bp and expected size range for miRNA targets is 75~200 bp.

<u>OPTION 2: real-time PCR</u>

1. Prepare the following reaction mix and load into a 384-well PCR plate.

| Ref | Components | Volume (μL) |
|---|---|---|
| Invitrogen S7563 | 50X SYBR Green | 0.1 |
| Invitrogen 12344040 | Accuprime Pfx SuperMix | 9.1 |
| | DP5 (20 μM) | 0.25 |
| | DP3 (20 μM) | 0.25 |
| | cDNA | 0.3 |

* Prepare triplicates for each sample, 1 μL cDNA is added to a master mix of 30 μL. Therefore, the optimal cycle number should be N-1. N is the optimal cycle determined by real-time PCR.

2. Monitor the amplification under a real-time thermocycler using the following program:

| Temperature | Time | |
|---|---|---|
| 95 °C | 2 min | |
| 95 °C | 20 s | 30 cycles |
| 58 °C | 30 s | |
| 68 °C | 20 s | |

3. Determine the highest amplification cycle N before the SYBR green signal reaches a plateau. The optimal amplification cycle, therefore, is N-1.

#### Step 13. Library preparation – pre-amplification of cDNA libraries

1. Amplify the cDNA library with the best cycle determined from previous diagnostic/real-time PCR. Prepare 6 PCR reactions for each library to be made using recipe described in **Step XII, OPTION1-1**.
2. Load each sample into one 15% TBE-Urea gel. Resolve the miRNA and target bands on gel. Stain the gel with SYBR gold nucleic acid gel stain.
3. Excise the miRNA band (~65 bp) and target (75~200 bp, usually a smear) separately from the gel.
4. Cut the bands into slices and place the gel pieces in a 0.5 mL eppendorf tube with a hole on the bottom.
5. Place the 0.5 mL tube in a 2 mL eppendorf tube. Pass the gel through the hole by centrifugation at 13,000 rpm, 4 °C for 1 min.

6. Weigh the gels and add 1-2 volumes of diffusion buffer.
7. Incubate the gel pieces with diffusion buffer in a thermomixer at 55 °C, 1,000 rpm for 30 min.
8. Centrifuge at 14,000 rpm, 4 °C for 1 min to clear the diffusion buffer.
9. Pass the supernatant through a Nanosep column (0.2 $\mu$m, PALL corporation ODM02C34).
10. Determined the volume of supernatant and add 3 volume of buffer QG (Qiagen MinElute Gel Extraction Kit 28606).
11. Pass the samples through the Qiagen MinElute spin columns. Wash two times with buffer PE.
12. Elute DNA with 10~20 $\mu$L $H_2O$

##### Step 14. Library construction – Introducing sequencing adaptors

1. Design library multiplexing strategy and assign different barcodes to samples to be run on the same lane in a illumina flow-cell.
2. Prepare the following PCR reaction mixtures:

| Ref | Components | Volume ($\mu$L) |
|---|---|---|
| Invitrogen 12344040 | Accuprime Pfx SuperMix | 27 |
| | DSFP (20 $\mu$M) | 0.5 |
| | DP3-Barcode (20 $\mu$M) | 0.5 |
| | Eluted DNA | 3 |

*Prepare three reactions for each library.

3. Amplify each library with different number of cycles using the following program.

| Temperature | Time | |
|---|---|---|
| 95 °C | 2 min | |
| 95 °C | 20 s | X cycles (x = 5, 7, 9…) |
| 58 °C | 30 s | |
| 68 °C | 40 s | |
| 68 °C | 5 min | |
| 4 °C | ∞ | |

4. Load the same library amplified by different cycles side-by-side onto a 6% TBE polyacrylamide gel (Invitrogen EC6265) and run the gel following standard protocol.
5. Select the best amplification cycle (usually the lowest) and cut the PCR product from gel.
6. Cut the gel into pieces and add 300 $\mu$L water to elute DNA. Incubate the gel with water on a rotator at 4 °C overnight.
7. Next day, remove gel pieces from water by passing it through a Nanosep column.

8. Precipitate DNA by adding 30 μL 3 M Sodium Acetate (pH5.5), 2 μL glycogen and 2 μL 0.1x NF-pellet paint (Novagen 70748-3) and 975 μL absolute Ethanol.
9. Pellet DNA by centrifugation at 4 °C, 14,000 rpm for 20 min.
10. Wash pellet with 500 μL 70% Ethanol.
11. Air-dry the pellet and resuspend in 15 μL water.
12. Submit the DNA libraries to the Integrated Genomics Operation Core at Memorial Sloan Kettering Cancer Center for quality-control, quantification, library pooling and high-throughput sequencing.

#### Input Control Library

##### Step 1. Dephosphorylation of RNA 3'ends

1. Prepare the following reaction mix:

| Ref | Components | Volume (μL) |
|---|---|---|
| Promega M183A | 10X Alkaline phosphatase buffer | 2.5 |
| Promega M182A | Alkaline phosphatase, Calf Intestine | 2.5 |
| Promega N251B | rRNasin | 0.5 |
| | Water | 9.5 |

2. Add the mix to 10 μL of lysates saved before Halo-Ago2 pulldown.
3. Incubate in a thermomixer at 37°C, 1,000 rpm for 20min.

##### Step 2. T4 PNK treatment

1. Prepare the following reaction mix:

| Ref | Components | Volume (μL) |
|---|---|---|
| NEB B0201S | 10X PNK buffer | 10 |
| NEB P0756S | ATP (10mM) | 1.25 |
| NEB M0201L | T4 PNK | 5 |
| Promega N251B | rRNasin | 2.5 |
| | Water | 56.25 |

2. Add 75 μL mix directly to each sample (25 μL).
3. Incubate in a thermomixer at 37°C, 1,000 rpm for 20min.

##### Step 3. RNA clean-up

Clean up input RNA by MyOne Silane Beads (Thermo 37002D) (Adapted from eCLIP protocol).

a) Prepare beads:

- Magnetically separate 20 $\mu$L MyONE Silane beads per sample, remove supernatant.
- Wash 1x with 900 $\mu$L RLT buffer (Qiagen 79216).
- Resuspend beads in 300 $\mu$L RLT buffer per sample.

b) Bind RNA:

- Add beads in 300 $\mu$L RLT buffer to each sample. Mix.
- Add 10 $\mu$L 5M NaCl and 615 $\mu$L Absolute Ethanol.
- Rotate at room temperature for 15 min.

c) Wash beads:

- Wash beads with 1 mL 75% Ethanol, pipette resuspend and move the suspension to a new tube.
- After 30 s, magnetically separate, remove supernatant.
- Wash beads 2x with 75% Ethanol.
- Briefly spin the tube in centrifuge, magnetically separate beads, remove supernatant.
- Air-dry the beads for 5 min.

d) Elute RNA:

- Resuspend beads in 10 $\mu$L H2O, let sit for 5 min.
- Magnetically separate.
- Transfer 10 $\mu$L of supernatant to new tube.

#### Step 4. 3' RNA linker ligation

1. Prepare the following reaction mix:

| Ref | Components | Volume ($\mu$L) |
|---|---|---|
| NEB B0216S | 10X T4 RNA ligase buffer | 2 |
| | BSA (0.2 $\mu$g/$\mu$l) | 2 |
| NEB P0756S | ATP (10 mM) | 2 |
| NEB M0204S | T4 RNA ligase 1 | 0.75 |
| Promega N251B | rRNasin | 0.5 |
| | RL3 (20 $\mu$M) | 1.25 |
| | Water | 1.5 |

2. Add 10 $\mu$L mix to each 10 $\mu$L of eluted RNAs.
3. Incubate in a thermomixer at 16°C overnight.

#### Step 5. RNA clean-up

Next day, clean up RNA using MyONE silane beads.

a) Prepare beads:

- Magnetically separate 20 $\mu$L MyONE Silane beads per sample, remove supernatant.

- Wash 1x with 900 μL RLT buffer (Qiagen 79216).
- Resuspend beads in 61.6 μL RLT buffer per sample.

b) Bind RNA:

- Add beads in 61.6 μL RLT buffer to each sample. Mix.
- Add 61.6 μL Absolute Ethanol.
- Incubate at room temperature for 15 min. Pipette mix every 3~5 min.

c) Wash beads:

- Wash beads with 1 mL 75% Ethanol, pipette resuspend and move the suspension to a new tube.
- After 30 s, magnetically separate, remove supernatant.
- Wash beads 2x with 75% Ethanol.
- Briefly spin the tube in centrifuge, magnetically separate beads, remove supernatant.
- Air-dry the beads for 5 min.

d) Elute RNA:

- Resuspend beads in ~6 μL H2O, let sit for 5 min.
- Magnetically separate.
- Transfer 5.9 μL of supernatant to new tube.

##### Step 6. 5' RNA adaptor ligation

(See Step 9 in HEAP library preparation starting from preparing 5' adaptor ligation mix)

##### Step 7. DNase treatment

(See Step 10 in HEAP library preparation)

##### Step 8. Reverse transcription for input RNA

(See Step 11 in HEAP library preparation)

##### Step 9. First PCR to determine optimal amplification cycles

Determine the optimal amplification conditions for input control libraries as instructed in Step 12 in HEAP library preparation. Based on experience, the optimal cycles are often between 13 to 18 cycles.

##### Step 10. Library preparation – pre-amplification of input cDNA libraries

Follow procedures in Step 13 in HEAP library preparation. However, a minor change is applied to the gel purification step. To prepare "size-matched" input control libraries for corresponding HEAP libraries, only extract PCR products between 75 and 200 bp from the 15% TBE-Urea gel.

#### Step 11. Library construction – Introducing sequencing adaptors

(See Step 14 in HEAP library preparation)

#### Appendix A: Buffer Recipes

**1x PBS**

| | |
|---|---|
| 137 mM | NaCl |
| 2.7 mM | KCl |
| 10 mM | $Na_2HPO_4$ |
| 1.8 mM | $KH_2PO_4$ |

**5x PBS**

| | |
|---|---|
| 685 mM | NaCl |
| 13.5mM | KCl |
| 50 mM | $Na_2HPO_4$ |
| 9 mM | $KH_2PO_4$ |

**1x TBS**

| | |
|---|---|
| 100 mM | Tris-HCl (pH7.5) |
| 150 mM | NaCl |

**Mammalian Lysis Buffer (Promega)**

| | |
|---|---|
| 50 mM | Tris-HCl (pH7.5) |
| 150 mM | NaCl |
| 1% | Triton X-100 |
| 0.1% | Na deoxycholate |

**Wash/Eq**

0.05% IGEPAL CA-630 in 1xTBS

**SDS Elution Buffer (10 mL)**

| | |
|---|---|
| 0.1% | SDS |
| 50 mM | Tris-HCl (pH7.5) |

**LiCl Wash Buffer**

| | |
|---|---|
| 100 mM | Tris-HCl (pH8.0) |
| 500 mM | LiCl |
| 1% | IGEPAL CA-630 |
| 1% | Na deoxycholate |

**PXL (1x)**

In 1X PBS, add:

| | |
|---|---|
| 0.1% | SDS |
| 0.5% | Na deoxycholate |
| 0.5% | IGEPAL CA-630 |

**PXL (5x)**

In 5X PBS, add:

| | |
|---|---|
| 0.1% | SDS |
| 0.5% | Na deoxycholate |
| 0.5% | IGEPAL CA-630 |

**1x PNK Buffer**

| | |
|---|---|
| 50 mM | Tris-HCl (pH7.4) |
| 10 mM | MgCl2 |
| 0.5% | IGEPAL CA-630 |

**1x PNK + EGTA**

| | |
|---|---|
| 50 mM | Tris-HCl (pH7.4) |
| 20 mM | EGTA |
| 0.5% | IGEPAL CA-630 |

**PK Buffer (Proteinase K)**

| | |
|---|---|
| 100 mM | Tris-HCl (pH7.5) |
| 50 mM | NaCl |
| 10 mM | EDTA |

**1x PK Buffer/7M Urea (prepare FRESH each time)**

| | |
|---|---|
| 100 mM | Tris-HCl (pH7.5) |
| 50 mM | NaCl |
| 10 mM | EDTA |
| 7M | Urea |

**Diffusion buffer**

| | |
|---|---|
| 0.5 M | Ammonium acetate |
| 10 mM | Magnesium acetate |
| 1 mM | EDTA (pH8.0) |
| 0.1% | SDS |

### Appendix B: Schematic of library construction

1. Linker ligation

RL5D-6 AGGGAGGACGAUGCGGNNNNNNNG RNA GUGUCAGUCACUUCCAGCGGpuro RL3

2. Reverse transcription

AGGGAGGACGAUGCGGNNNNNNNG RNA GUGUCAGUCACUUCCAGCGGpuro
CACAGTCAGTGAAGGTCGCC DP3

3. Amplification

DP5 AGGGAGGACGATGCGG
AGGGAGGACGATGCGGNNNNNNNG DNA GTGTCAGTCACTTCCAGCGG
TCCCTCCTGCTACGCCNNNNNNNC DNA CACAGTCAGTGAAGGTCGCC
CACAGTCAGTGAAGGTCGCC DP3

4. Library construction

DSFP5
AGGGAGGACGATGCGGNNNNNNNG DNA GTGTCAGTCACTTCCAGCGG
TCCCTCCTGCTACGCCNNNNNNNC DNA CACAGTCAGTGAAGGTCGCC
DP3-Barcode

DSFP5:

AATGATACGGCGACCACCGACTATGGATACTTAGTCAGGGAGGACGATGCGG

DP3-Barcode:

CAAGCAGAAGACGGCATACGAGATNNNNNNNNGTGACTGGAGTTCAGACGTGTGCTCTTCCGATCCCGCTGGAAGTGACTGACAC
Illumina TruSeq index

5. Sequencing strategy

Illumina TruSeq index read primer SP2

AATGATACGGCGACCACCGACTATGGATACTTAGTCAGGGAGGACGATGCGGNNNNNNNG DNA GTGTCAGTCACTTCCAGCGGGATCGGAAGAGCACACGTCTGAACTCCAGTCACNNNNNNNNATCTCGTATGCCGTCTTCTGCTTG
TTACTATGCCGCTGGTGGCTGATACCTATGAATCAGTCCCTCCTGCTACGCCNNNNNNNC DNA CACAGTCAGTGAAGGTCGCCCTAGCCTTCTCGTGTGCAGACTTGAGGTCAGTGNNNNNNNNTAGAGCATACGGCAGAAGACGAAC
CTATGGATACTTAGTCAGGGAGGACGATGCGG
Customized sequencing primer SSP1
