## Supplementary Figures 1-14 for "High-resolution *in vivo* identification of miRNA targets by Halo-Enhanced Ago2 Pulldown"

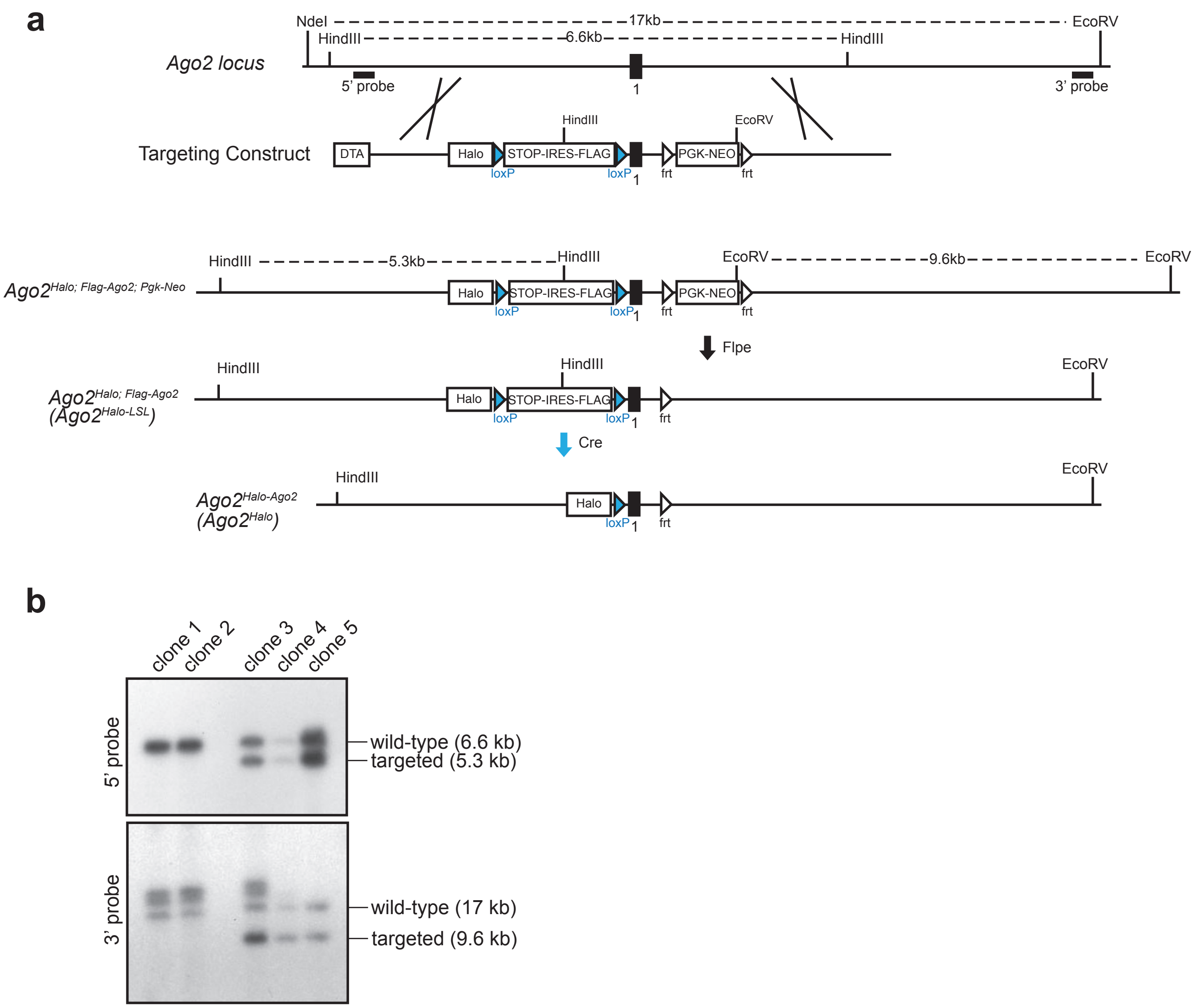


**Figure S1. Generation of the Halo-Ago2 conditional allele.**
**a)** Targeting strategy for the generation of conditional Halo-Ago2 knock-in mESCs . Halo, HaloTag; DTA, diphtheria toxin A; STOP, stop codon; IRES, internal ribosome entry site; frt, flippase recognition target; Flpe, flippase recombinase. **b)** mESC single clones were screened by southern blot using probes mapping to sequences outside each of the 5' and 3' homology regions as shown in **a).**

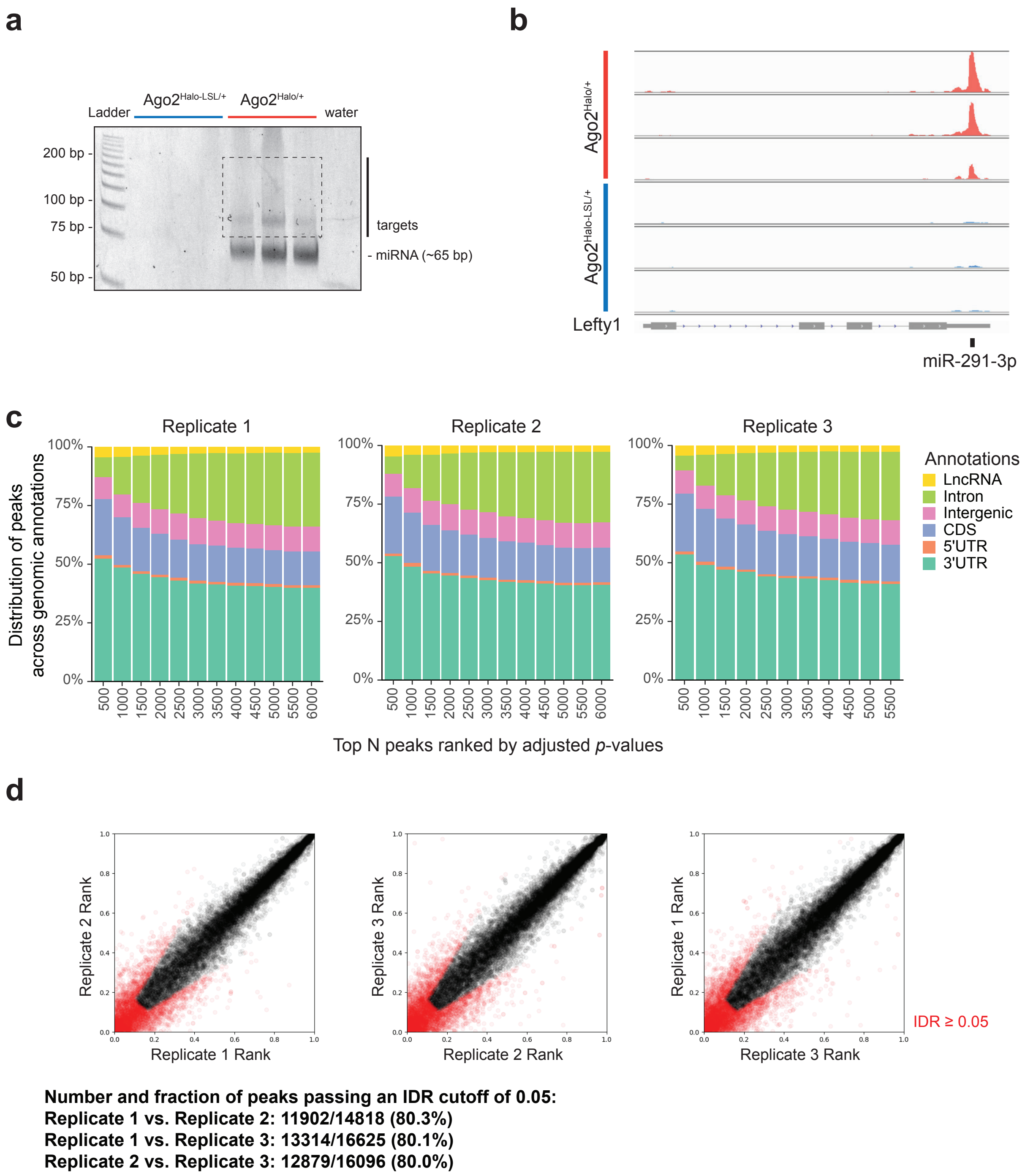


**Figure S2 Generation of HEAP libraries from mESCs.**
**a)** Polyacrylamide gel electrophoresis of PCR products of HEAP libraries generated from Ago2$^{Halo-LSL/+}$ and Ago2$^{Halo/+}$ mESCs. After reverse transcription, cDNAs were amplifed for 17 cycles. Amplification products were only seen in libraries from mESCs expressing the Halo-Ago2 fusion. The PCR generated a band corresponding to miRNA (~65 bp) and a smear corresponding to miRNA targets (75~200 bp). **b)** Genome browser view of the Lefty1 3'UTR with tracks corresponding to HEAP mRNA libraries generated from Ago2$^{Halo/+}$ (N = 3) and Ago2$^{Halo-LSL/+}$ (N = 3) mESCs. **c)** Peaks identified in each HEAP library replicate were ranked by increasing adjusted *p*-value before calculating their distribution across genomic annotations. **d)** Pairwise IDR analysis between HEAP mESC biological replicates. Each point represents the rank of an individual peak as determined in each pair of the replicates. Points in red correspond to peaks that failed to pass an IDR cutoff of 0.05. Table at the bottom summarizes the absolute number and percentage of peaks that passed the 0.05 IDR cutoff in each pair of the comparison.

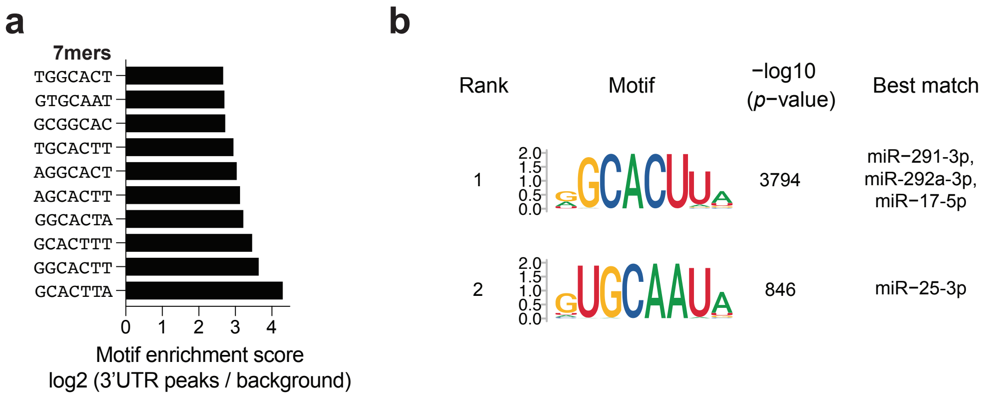


**Figure S3 Motif discovery in HEAP libraries from mESCs.**

**a)** Bar plot showing the top ten 7-mers enriched in Halo-Ago2 3'UTR binding sites as determined by unbiased motif enrichment analysis. Motif enrichment score was calculated as log2 [($Freq_{3'UTR}$ + c) / ($Freq_{bg}$ + c)]. $Freq_{3'UTR}$: frequencies of 7-mers in 3'UTR peaks. $Freg_{bg}$: frequencies of 7-mers in background sequences, which were defined as 3'UTR sequences outside of peaks. **b)** Sequence logos of the most enriched 8-mer motifs in HEAP 3'UTR peaks as determined by the HOMER *de novo* motif discovery algorithm. miRNA families whose seeds are complementary to these motifs are shown.

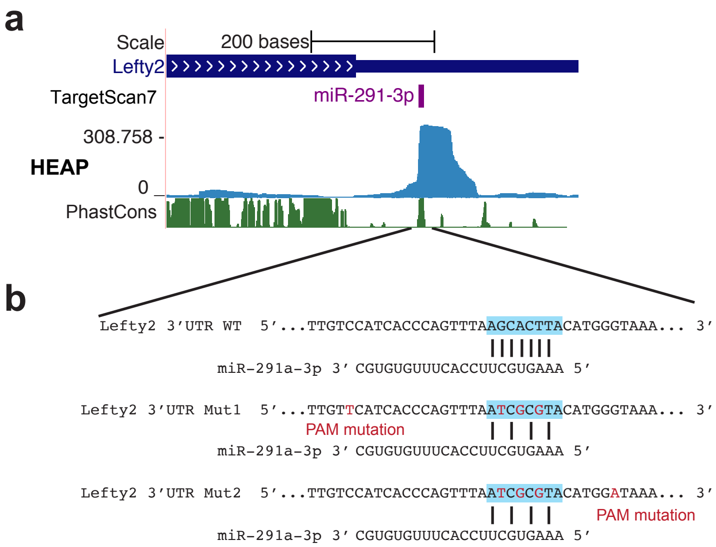


**Figure S4. Generation of the Lefty2 3'UTR mutant mESCs.**
**a)** Genome browser view of the 3'UTR peak identified in Lefty2. The miR-291-3p binding site predicted by TargetScan7 is annotated. PhastCons: placental mammal conservation by PhastCons. **b)** Around the predicted miR-291-3p binding site in the *wild-type* and in the *Lefty2*$^{MUT}$ mESCs. The mutations introduced are highlighted in red and disrupt the base-pairing between Lefty2 3'UTR and miR-291a-3p. PAM sequence mutations specific to each of the guide RNAs used are also highlighted.

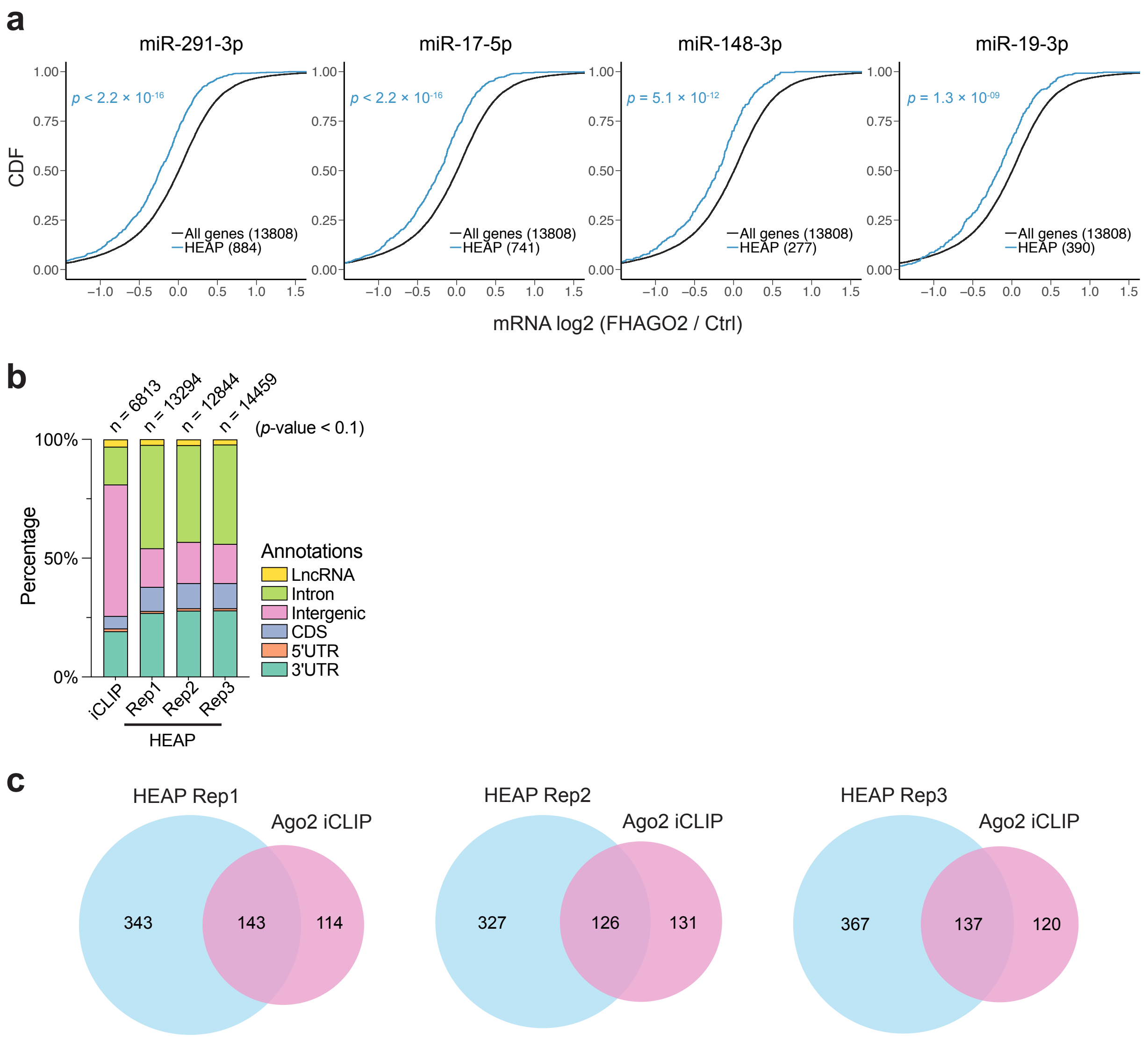


**Figure S5 Assessing functionality of Halo-Ago2 binding sites and the comparison between iCLIP and HEAP.**
**a)** Cumulative distribution function (CDF) plots for targets of selected miRNA families identified by HEAP. The mRNA log2 fold change was calculated in Ago1-4$^{-/-}$ mESCs upon ectopic FHAGO2 expression (Bosson et al., 2014; GSE61348; *p*-value: one-sided Kolmogorov–Smirnov test). **b)** Genomic distribution of Ago2 binding sites identified in the iCLIP library and in three HEAP libraries at a *p*-value cutoff of 0.1. The total number of peaks identified in each library is shown. **c)** Venn diagrams showing the overlap between miR-291-3p 3'UTR binding sites identified by iCLIP and those identified by each HEAP library.

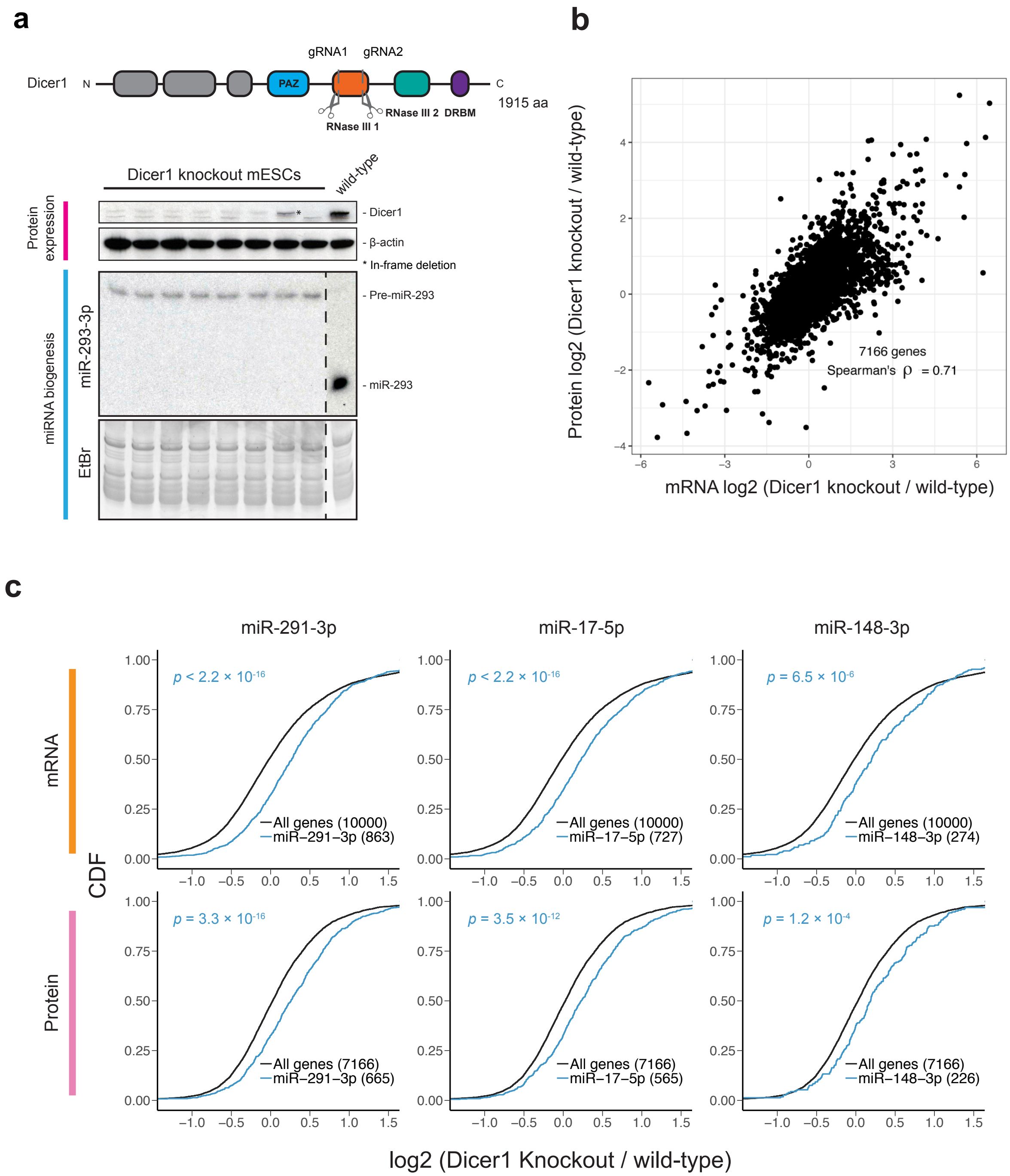


**Figure S6. Assessing HEAP-identified miRNA binding sites in Dicer1 knockout mESCs.**
**a)** Generation of Dicer1 knockout mESCs using CRISPR-cas9. Schematic of the Dicer1 protein is shown in the upper panel, with key domains annotated. A portion of the first RNase III domain was deleted using a pair of guide RNAs targeting the corresponding location on the genome. Dicer1 knockout single clones were identified using immunoblot and northern blot. Notice the complete loss of Dicer1 full-length protein and the blockade of Pre-miR-293 maturation in the Dicer1 knockout mESC clones. DRBM: double-stranded RNA binding motif. Vertical dashed line indicates splicing of the Northern membrane needed to align RNA and protein samples. **b)** Changes in mRNA expression between the Dicer1 knockout and wild-type mESCs highly correlate with changes in protein expression. mRNA expressions were measured by RNA-seq. Protein expressions were measured using mass spectrometry proteomics. **c)** Cumulative distribution function (CDF) plots for targets of miR-291-3p, miR-17-5p and miR-148-3p identified by HEAP. The log2 fold changes in mRNA and protein levels were calculated as Dicer1 knockout vs. wild-type mESCs (*p*-value: one-sided Kolmogorov–Smirnov test).

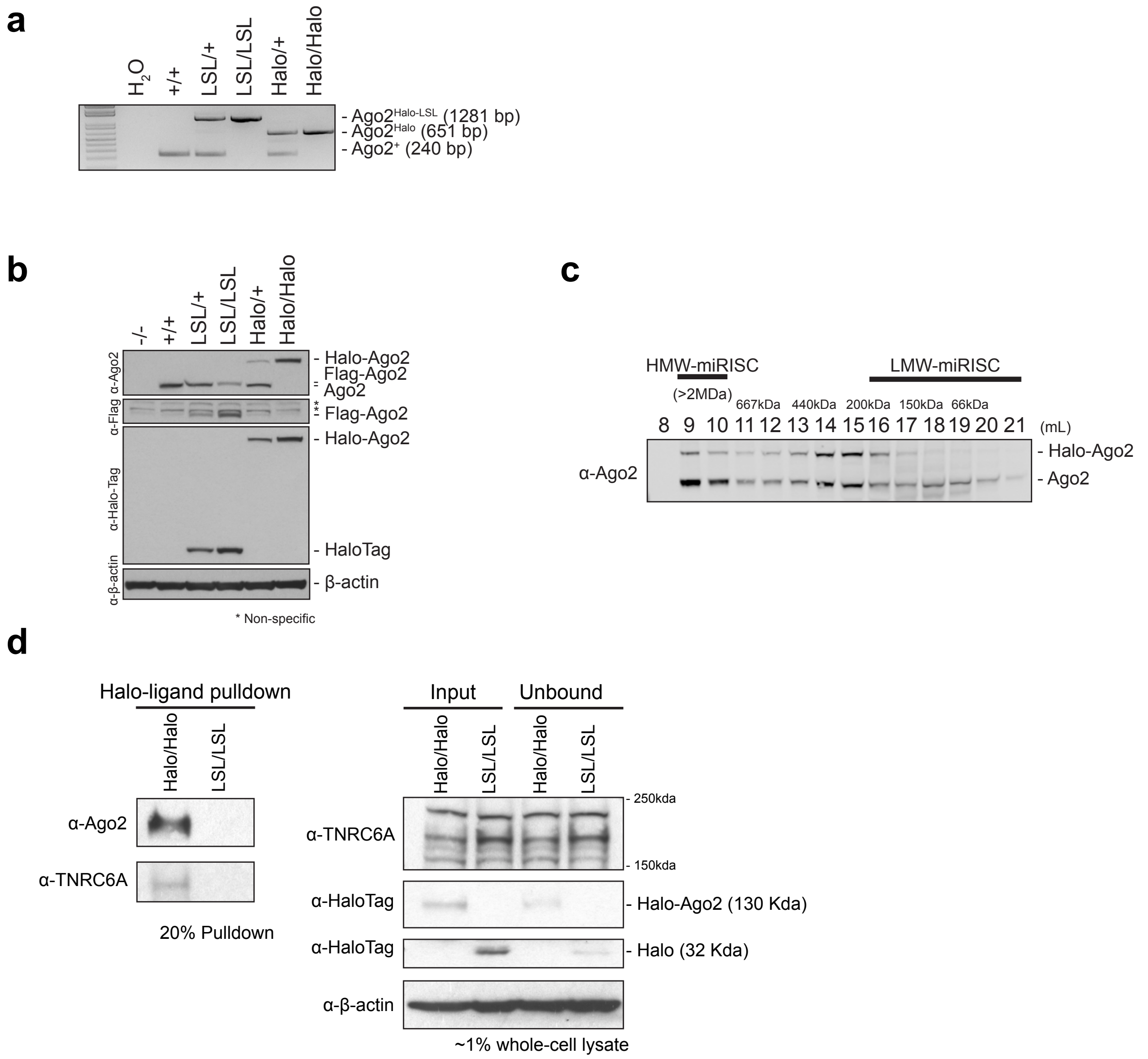


**Figure S7 Expression of Halo-Ago2 and its interaction with TNRC6A**

**a)** Genotyping PCR of MEFs derived from E13.5 embryos harboring the indicated alleles. +: the wild-type allele or Ago2$^{+}$; LSL: Ago2$^{Halo\text{-}LSL}$; Halo: Ago2$^{Halo}$. **b)** Whole-cell lysates of MEFs of indicated genotypes were probed with antibodies against Ago2, Flag, HaloTag and β-actin. **c)** Size exclusion chromatography fractionated Ago2$^{Halo/+}$ MEF lysates were probed with an antibody against Ago2. HMW: high-molecular weight; LMW: low-molecular weight; miRISC: miRNA-induced silencing complex. **d)** The HaloTag was pulled down from whole-cell lysates of Ago2$^{Halo\text{-}LSL/Halo\text{-}LSL}$ (LSL/LSL) and Ago2$^{Halo/Halo}$ (Halo/Halo) MEFs using HaloTag ligand conjugated to magnetic beads. Proteins co-purified were probed with antibodies against Ago2 and TNRC6A. Input and unbound lysates were probed with antibodies against TNRC6A, HaloTag and β-actin.

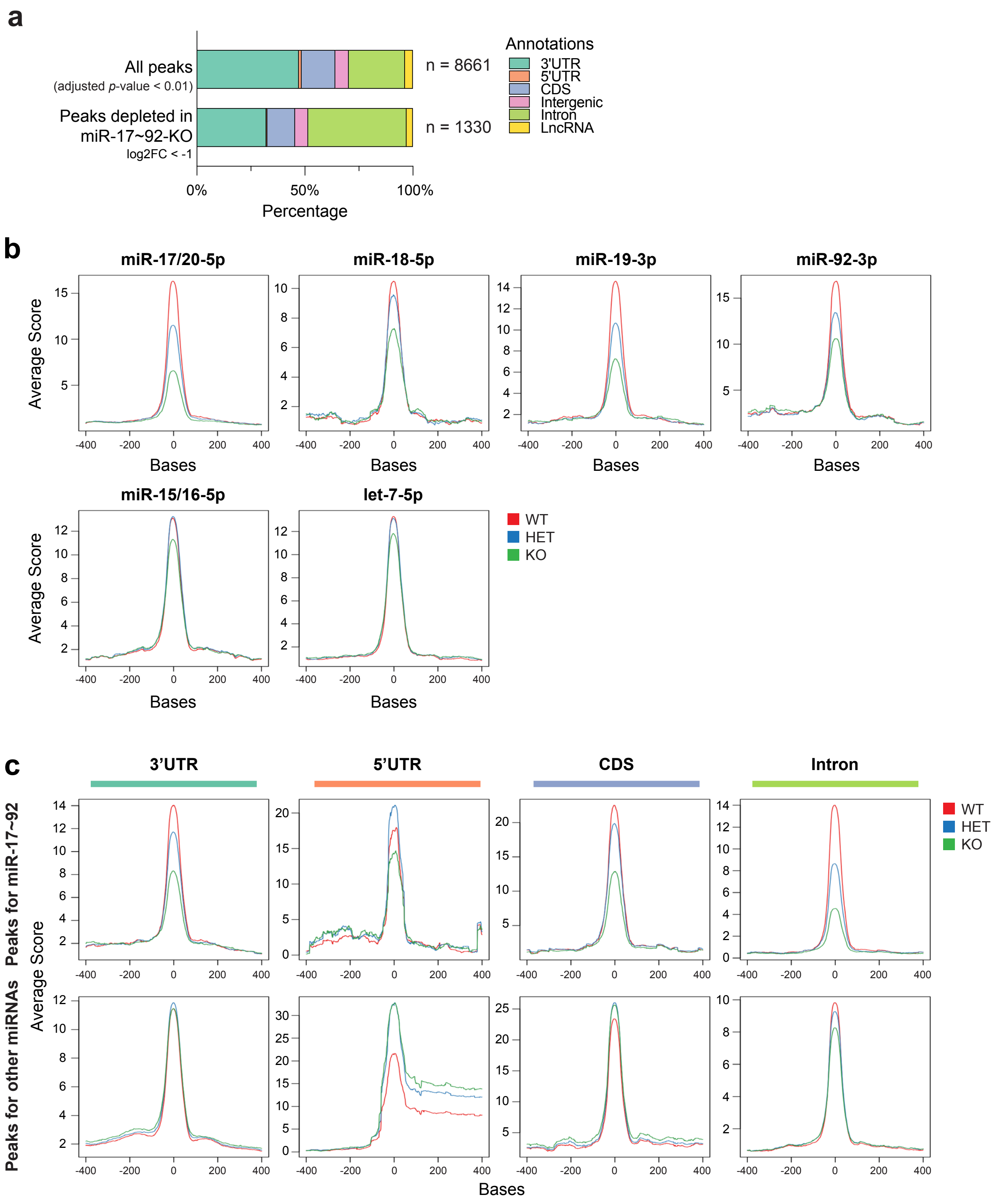


**Figure S8 Identification of miRNA binding sites in wild-type and miR-17~92-null embryos.**

**a)** Bar charts showing the number and distribution across genomic annotations of peaks identified in the HEAP libraries from E13.5 embryos. "All peaks" indicates peaks identified across all genotypes. Peaks selectively depleted in the miR-17-92-KO embryo are shown in the lower panel. **b-c)** Histogram of average score of read counts in an 800-bp region surrounding HEAP peaks in miR-17~92-WT, miR-17~92-HET and miR-17~92-KO E13.5 embryos. In **b)**, peaks containing seed matches for miRNAs belonging to the miR-17~92 cluster are plotted in the upper panels, while peaks containing seed matches for two other miRNA families are plotted in the lower panels. In **c)**, peaks containing seed matches for the top 31 miRNA families ranked by abundance were chosen and grouped by their genomic locations. Peaks containing seed matches for miRNAs belonging to the miR-17~92 cluster are plotted in the upper panels, while the remaining peaks are plotted in the lower panels.

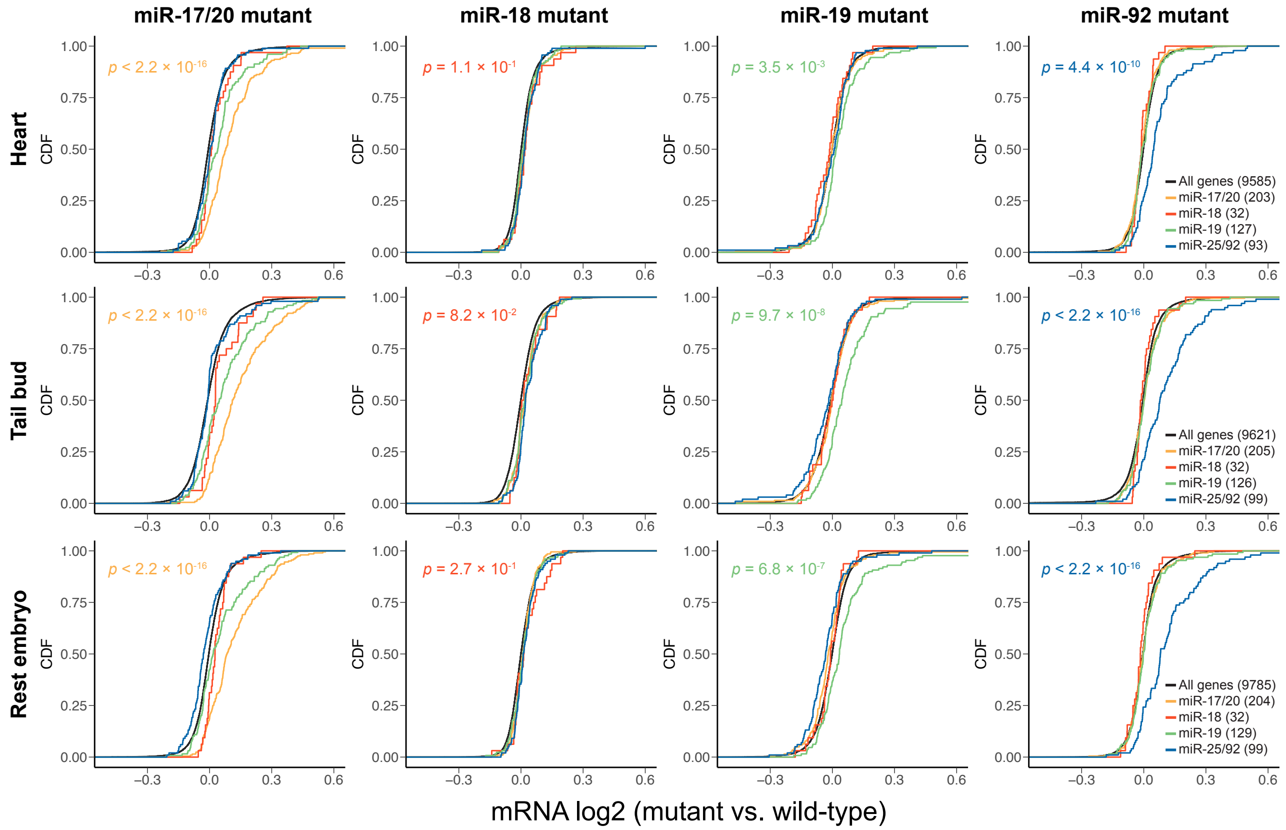


**Figure S9 Assessing functionality of HEAP targets identified in E13.5 embryos.**
Cumulative distribution function (CDF) plots for HEAP targets containing seed matches for the miR-17/20-5p, miR-18-5p, miR-19-3p and miR-25/92-3p seed families. The mRNA log2 fold change for a particular miRNA (e.g. miR-17/20) was calculated in the miR-17~92 mutant E9.5 embryos (Han et al., 2015; GSE63813) by contrasting all conditions that are mutant for this miRNA (e.g. Δ17, Δ17,18, Δ17,18,92 and KO) to all other conditions (e.g. Δ18, Δ19, Δ92). KO: embryos null for the entire miR-17~92 cluster; Δ17: embryos null for miR-17 and miR-20a; Δ18: embryos null for miR-18a; Δ19: embryos null for miR-19a and miR-19b-1, Δ92: embryos null for miR-92a-1; Δ17,18: embryos null for miR-17, miR-18a and miR-20a; Δ17,18,92: embryos null for miR-17, miR-18a, miR-20a and miR-92a-1. *P*-values were calculated between "all genes" and targets of the corresponding deleted miRNA seed families using one-sided Kolmogorov–Smirnov tests.

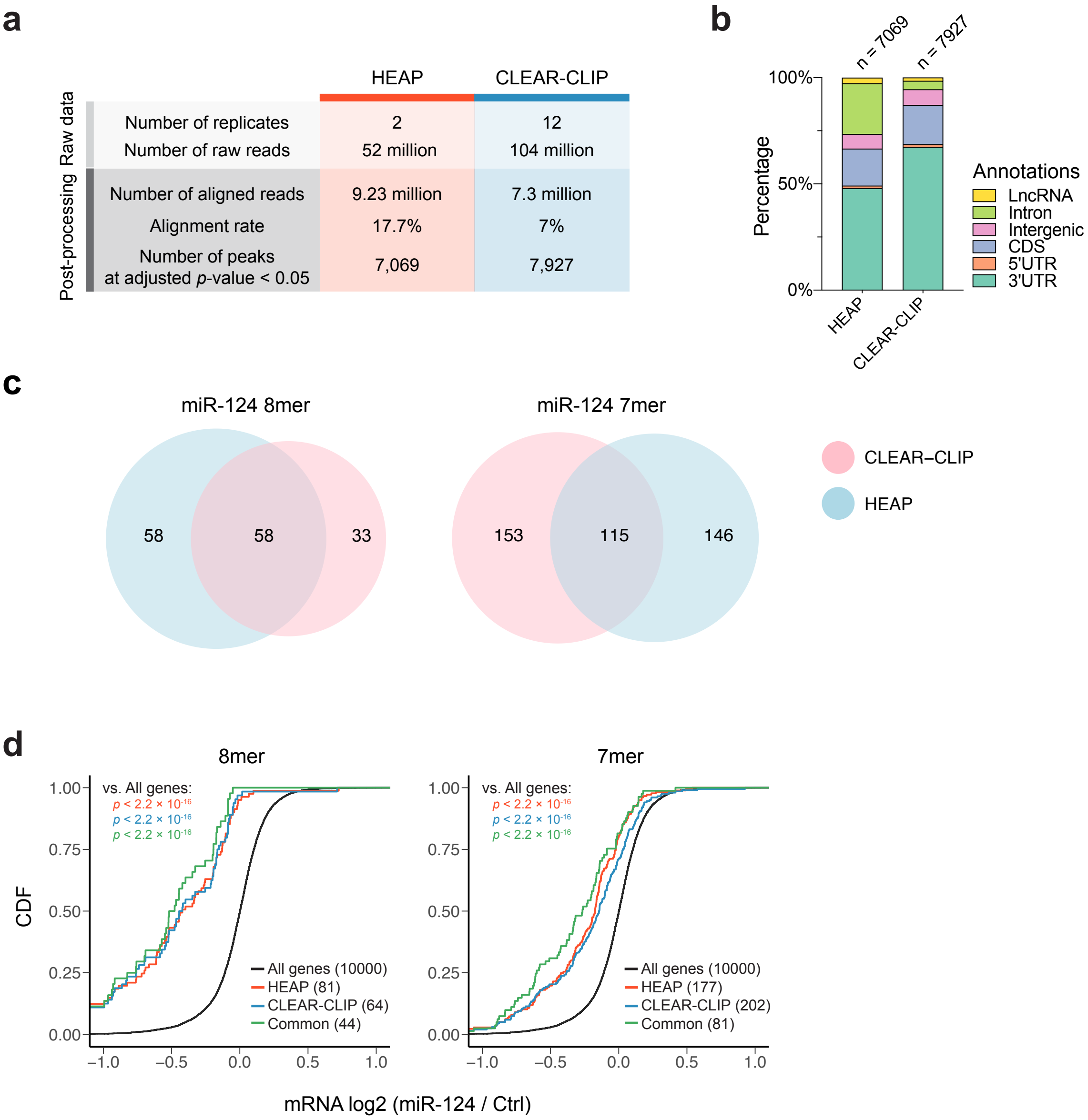


**Figure S10 Comparison between HEAP and CLEAR-CLIP.**
**a)** Comparison between HEAP and CLEAR-CLIP libraries generated from cortices of P13 mice (Moore et al. 2015; GSE73059). Peaks were filtered using an adjusted *p*-value cutoff of 0.05. **b)** Bar plot showing genomic distribution of peaks identified by HEAP and CLEAR-CLIP. **c)** Venn diagrams showing the overlap between miR-124-3p target genes identified by CLEAR-CLIP and those identified by HEAP. Targets were grouped by seed matches types. **d)** Cumulative distribution function (CDF) plots for targets of miR-124-3p identified by HEAP, by CLEAR-CLIP or by both methods (common). Targets with 8mer (left) or 7mer (right) seed matches for miR-124-3p were plotted separately. mRNA log2 fold changes were obtained from a dataset generated from a mouse neuroblastoma cell line (CAD) overexpressing miR-124 (Mekeyev et al., 2007; GSE8498; *p*-value: one-side Kolmogorov-Smirnov test).

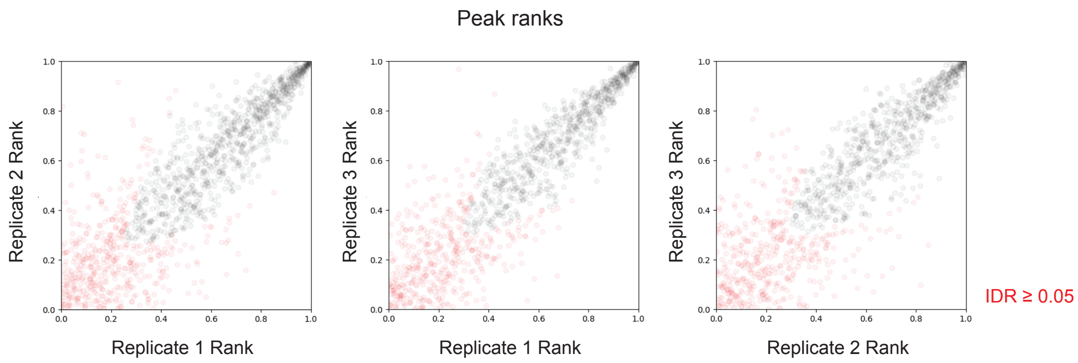


**Number and fraction of peaks passing an IDR cutoff of 0.05:**
**Replicate 1 vs. Replicate 2: 775/1235 (62.8%)**
**Replicate 1 vs. Replicate 3: 627/1058 (59.3%)**
**Replicate 2 vs. Replicate 3: 628/1054 (59.6%)**

**Figure S11 Reproducibility of HEAP libraries from Bcan-Ntrk1 gliomas.**
Pairwise IDR analysis between biological replicates of HEAP libraries generated from the Bcan-Ntrk1 gliomas. Each point represents the rank of an individual peak as determined in each pair of the replicates. Points in red correspond to peaks that fail to pass an IDR cutoff of 0.05. Table at the bottom summarizes the absolute number and percentage of peaks that passed the 0.05 IDR cutoff in each pair of the comparison.

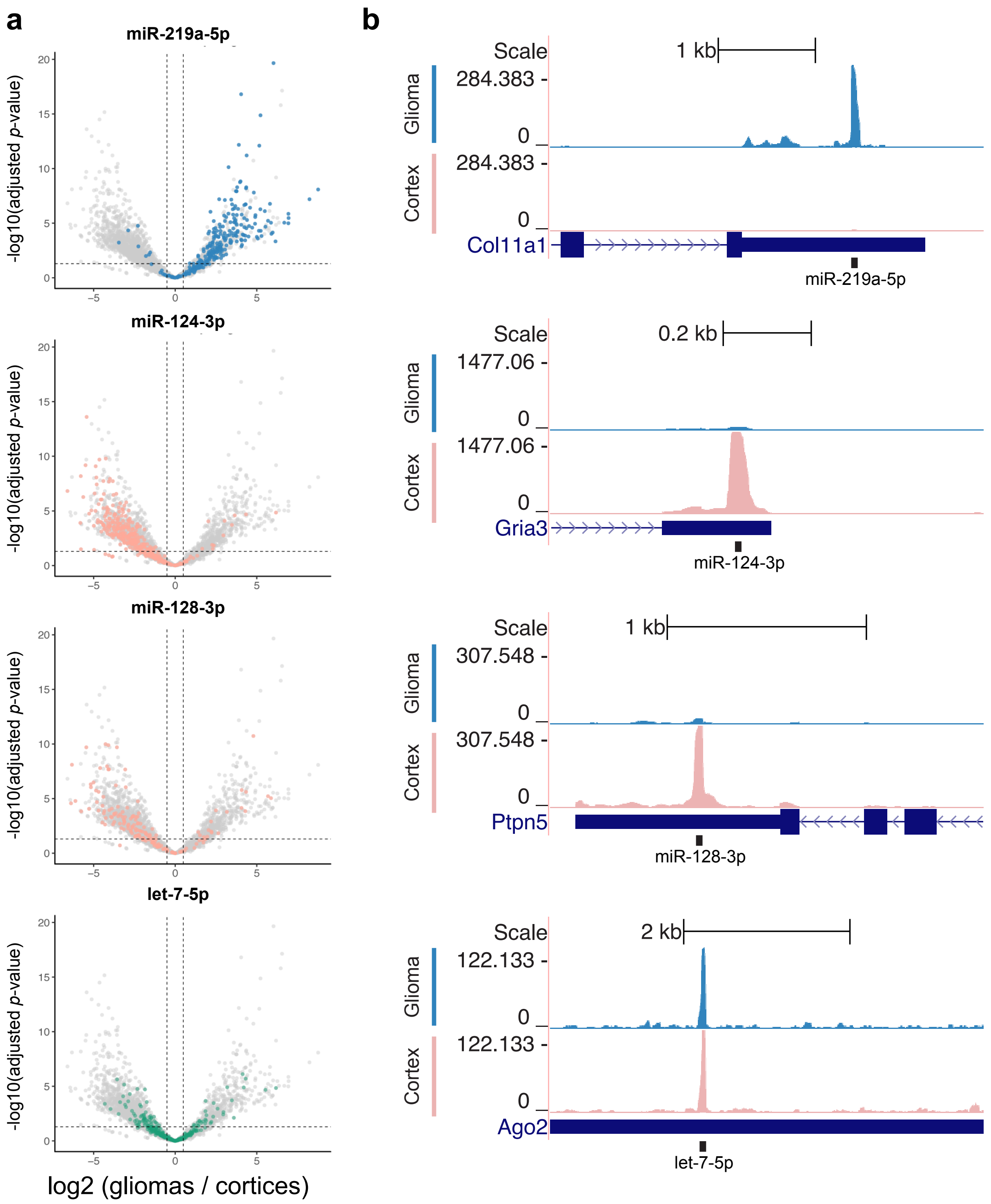


**Figure S12 miRNA binding sites in Bcan-Ntrk1 gliomas and cortices.**
**a)** Volcano plots of changes in peaks signal intensities between Bcan-Ntrk1 gliomas and cortices. Peaks containing seed matches for selected miRNA families are highlighted. **b)** Genome browser view of representative peaks for miRNA families in **a)**.

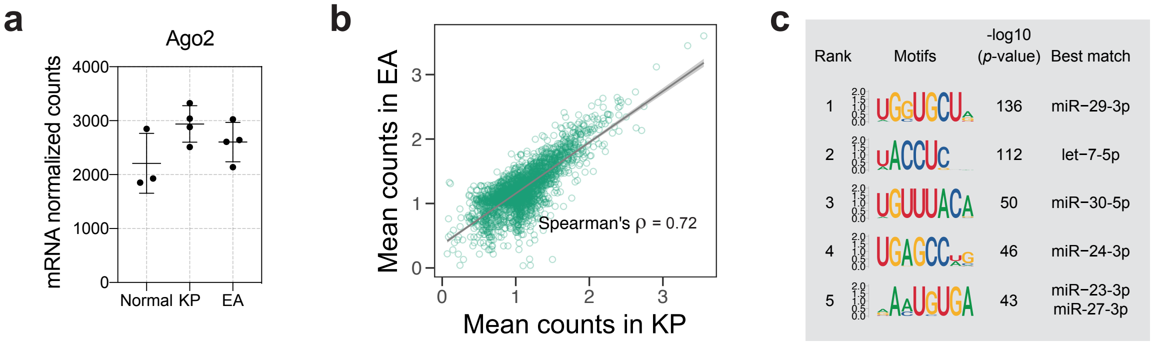


**Figure S13 HEAP in lung and lung adenocarcinomas.**

**a)** Normalized counts for Ago2 mRNA in normal lung and in KP and EA lung tumors. **b)** Scatter plot for normalized peak counts as detected in KP (x-axis) or EA (y-axis) HEAP libraries. **c)** Sequence logos of the most enriched 8-mers as determined by the HOMER *de novo* motif discovery algorithm under HEAP peaks identified in normal lungs (cutoff: adjusted *p*-value < 0.05; log2 (lungs/ tumors) > 0.5). miRNA families whose seeds are complementary to these motifs are annotated .

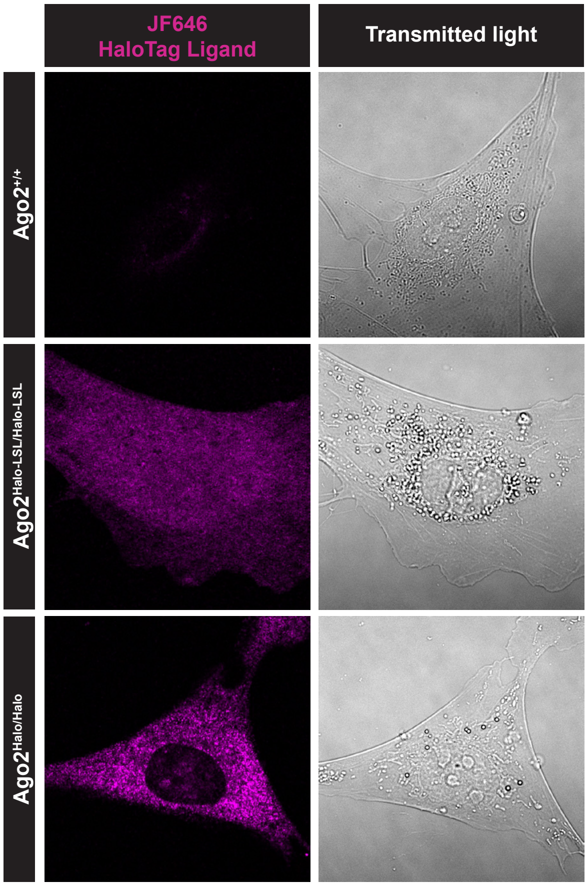


**Figure S14 Live-cell imaging of endogenous Halo-Ago2.**
Confocal imaging of MEFs of indicated genotypes. MEFs were incubated with 200 mM Janelia Fluor 646 HaloTag ligand 1 hr pior to the imaging.
